# Deconvolution of Expression for Nascent RNA Sequencing Data (DENR) Highlights Pre-RNA Isoform Diversity in Human Cells

**DOI:** 10.1101/2021.03.16.435537

**Authors:** Yixin Zhao, Noah Dukler, Gilad Barshad, Shushan Toneyan, Charles G. Danko, Adam Siepel

## Abstract

Quantification of mature-RNA isoform abundance from RNA-seq data has been extensively studied, but much less attention has been devoted to quantifying the abundance of distinct precursor RNAs based on nascent RNA sequencing data. Here we address this problem with a new computational method called Deconvolution of Expression for Nascent RNA sequencing data (DENR). DENR models the nascent RNA read counts at each locus as a mixture of user-provided isoforms. The performance of the baseline algorithm is enhanced by the use of machine-learning predictions of transcription start sites (TSSs) and an adjustment for the typical “shape profile” of read counts along a transcription unit. We show using simulated data that DENR clearly outperforms simple read-count-based methods for estimating the abundances of both whole genes and isoforms. By applying DENR to previously published PRO-seq data from K562 and CD4^+^ T cells, we find that transcription of multiple isoforms per gene is widespread, and the dominant isoform frequently makes use of an internal TSS. We also identify > 200 genes whose dominant isoforms make use of different TSSs in these two cell types. Finally, we apply DENR and StringTie to newly generated PRO-seq and RNA-seq data, respectively, for human CD4^+^ T cells and CD14^+^ monocytes, and show that entropy at the pre-RNA level makes a disproportionate contribution to overall isoform diversity, especially across cell types. Altogether, DENR is the first computational tool to enable abundance quantification of pre-RNA isoforms based on nascent RNA sequencing data, and it reveals high levels of pre-RNA isoform diversity in human cells.

## Introduction

For about the last 15 years, most large-scale transcriptomic analyses have relied on high-throughput short-read sequencing technologies as the readout for the relative abundances of RNA transcripts. In species with available genome assemblies, these sequence reads are generally mapped to assembled contigs, and then the “read depth,” or average density of aligned reads’ is used as a proxy for the abundance of RNAs corresponding to each annotated transcription unit. The approach is relatively inexpensive and straightforward, and, with adequate sequencing depth, it generally leads to accurate estimates of abundance.

A fundamental challenge with this general paradigm, however, is that transcription units frequently overlap in genomic coordinates—that is, the same segment of DNA often serves as a template for multiple distinct RNA transcripts. As a result, it is unclear which transcription unit is the source of each sequence read. While this problem can occur at the level of whole genes that contain overlapping segments, it is most prevalent at the level of multiple isoforms for each gene, owing to alternative transcription start sites (TSSs), alternative transcription termination sites (TTSs) or polyadenylation and cleavage sites (PAS), and alternative splicing. These isoforms often overlap heavily with one another, and differ on a scale that is not well described by short-read sequencing. For example, a typical Illumina RNA sequencing run today generates reads of length 150 bp, roughly the size of an exon in the human genome. As a result, many reads fall within a single exon and therefore carry no direct information about the relative abundances of isoforms containing that exon. This problem is critical because the existence of multiple isoforms per gene is the rule rather than the exception in most eukaryotes. For example, more than 90% of multi-exon human genes undergo alternative splicing (1), with an average of more than 7 isoforms per protein-coding gene (2); in plants, up to 70% of multi-exon genes show evidence of alternative splicing (3).

In the case of RNA-seq data, the problem of isoform abundance estimation from short-read sequence data has been widely studied for more than a decade (4–6). Several software packages now address the problem efficiently and effectively, including ones that make use of fully mapped reads (7–10) and others that substantially boost speed by working only with “pseudoalignments” at remarkably little (if any) cost in accuracy (11–13). These computational methods differ in detail but they generally work by modeling the observed sequence reads as an unknown mixture of isoforms at each locus. They estimate the relative abundances (mixture coefficients) of the isoforms from the read counts, relying in particular on the subset of reads that reflect distinguishing features, such as exons or splice junctions present in some isoforms but not others. Because RNA-seq libraries are typically dominated by mature RNAs, intronic reads tend to be rare and splice junctions provide one of the strongest signals for differentiation of isoforms. Altogether, these isoform quantification methods work quite well, with the best methods exhibiting Pearson correlation coefficients of 0.95 or higher with true values in simulation experiments, and similarly high concordance across technical replicates for real data (2).

In recent years, another method for interrogating the transcriptome, known as “nascent RNA sequencing,” has become increasingly widely used. Instead of measuring the concentrations of mature RNAs, as RNA-seq effectively does, nascent RNA sequencing protocols isolate and sequence newly transcribed RNA segments, typically by tagging them with selectable ribonucleotide analogs or through isolation of polymerase-associated RNAs (14–22). In this way, they provide a measurement of primary transcription, independent of the RNA decay processes that influence cellular concentrations of mature RNAs. In addition, nascent RNA sequencing methods have a wide variety of other applications, including identification of active enhancers (through the presence of eRNAs) (20,22–25), characterization of promoter-proximal pausing and divergent transcription (14,15), estimation of elongation rates (26,27), and estimation of relative RNA half-lives (28).

In nascent RNA sequencing, the isolated RNAs have generally not yet been spliced; therefore, they represent the entire transcribed portion of the genome, including introns. As a result, the problem of distinguishing alternative splice forms is largely irrelevant. On the other hand, the data typically still reflect a mixture of precursor RNA (pre-RNA) isoforms, having different TSSs and/or TTSs/PASs. Moreover, the problem of decomposing this mixture can be more challenging than for RNA-seq in some respects, both because pre-RNA isoforms have fewer differentiating features than mature RNA isoforms, and because nascent RNA read depths tend to be substantially reduced, since introns as well as exons are sequenced. Distinguishing among pre-RNA isoforms in nascent RNA sequence data can be critical for a wide variety of downstream analyses. Nevertheless, to our knowledge, only one computational tool has been developed to address this problem—a program called TuSelector that was introduced in ref. (25)—and it has never been packaged for use by other research groups or rigorously evaluated for accuracy. In most analyses of nascent RNA sequencing data, the isoform deconvolution problem is either ignored or addressed by simple heuristics, such as assuming each gene is represented by the longest annotated isoform (e.g., refs. (29,30)).

In this article, we introduce a new computational method and implementation in R, called Deconvolution of Expression for Nascent RNA sequencing (DENR), that addresses the problem of isoform abundance quantification at the pre-RNA level. DENR also solves the closely related problems of estimating abundance at the gene level, summing over all isoforms, and identifying the “dominant isoform,” that is, the one exhibiting the greatest abundance. DENR makes use of a straightforward non-negative least-squares strategy for decomposing the mixture of isoforms present in the data, but then improves on this baseline approach by taking advantage of machine-learning predictions of TSSs and an adjustment for the typical shape profile in the read counts along a transcription unit. We show that the method performs well on simulated data, and then use it to reveal a high level of diversity in the pre-RNA isoforms inferred from PRO-seq data for several human cell types, including K562, CD4^+^ T cells, and CD14^+^ monocytes.

## Results

### Overview of DENR

DENR is implemented as a package in the R programming environment. It requires two main inputs: a set of isoform annotations and a set of corresponding strand-specific nascent RNA sequencing read counts. Mature RNA isoform annotations can be easily downloaded by making use of biomaRt (31) or extracted from files in commonly available formats, such as GTF or GFF; similarly read counts can be obtained from a file in bigWig format. Detailed examples are provided in an online vignette (see **Methods**).

Given the necessary inputs, DENR first builds a *transcript_quantifier* object, which summarizes the read counts corresponding to the available isoform annotations (**Fig. 1**). This phase consists of three steps (**Supplementary Fig. S1**). First, the mature RNA isoforms are grouped into nonoverlapping, strand-specific clusters, corresponding roughly to genes (although if two genes overlap on the same strand, they will be grouped in the same cluster). Second, masking rules are applied to the read counts, causing a user-specified number of bins to be excluded at the start and end of each annotated isoform, to avoid the biases in quantification stemming from promoter-proximal pausing or termination-related deceleration of RNA polymerase. Throughout this paper, we assume a bin size of 250 bp. Third, the set of mature isoforms in each cluster is collapsed to a maximal set such that each isoform model has a unique pair of start and end coordinates, by merging all mature isoforms that share both their start and end bins. This step reduces isoforms annotated at the mature RNA level, many of which differ only in their splice patterns, to a more compact set of pre-RNA isoforms. It also merges pre-RNA isoforms that differ from one another after masking. This second property is useful because the nascent RNA sequence data typically provides only approximate indications of the TSS and TTS associated with each transcript, owing to both sparseness of the data and imprecisions in the transcription process itself (such as transcriptional run-on at the 3’end). The reduced set represents isoforms likely to be confidently distinguishable on the basis of nascent RNA sequence data alone.

**Fig. 1.**
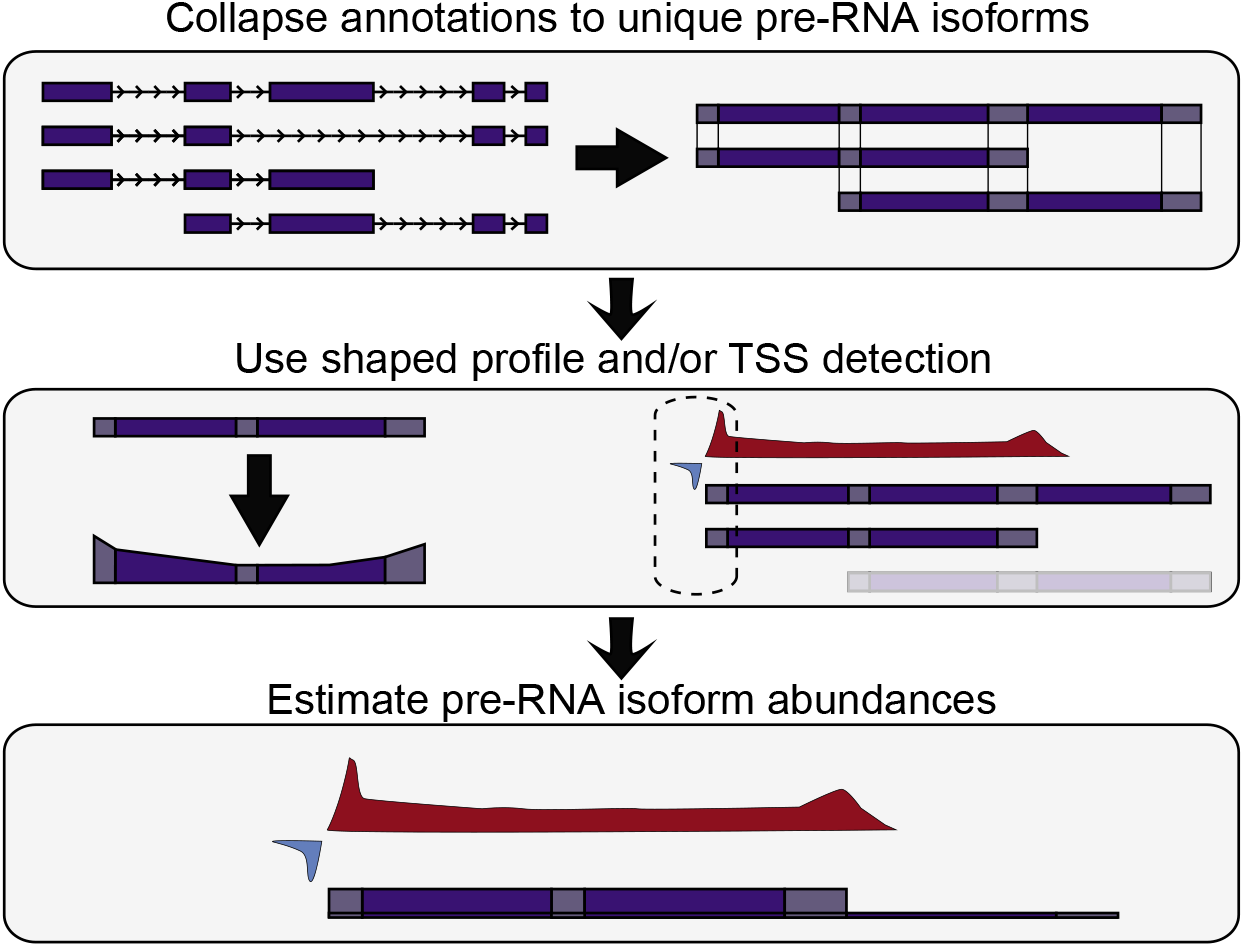
Illustration of DENR analysis. (*Top*) DENR first groups the available isoform annotations into nonoverlapping, stand-specific clusters and summarizes the associated read counts in genomic bins of user-specified size (default 250 bp). At this stage, it optionally masks bins corresponding to the start and end of each isoform. It then collapses mature RNA isoforms together that share start (TSS) and end (TTS/PAS) coordinates within the resolution of a single bin. (*Middle*) The program then optionally adjusts the isoform model to reflect a typical “U”-shaped profile, and optionally applies a machine-learning method to predict active TSSs based on patterns of bidirectional transcription. At this stage, it may also exclude isoforms designated by the user as inactive (not shown). (*Bottom*) Finally, DENR estimates the abundance of each isoform in each cluster by minimizing the squared difference between the expected and observed read counts across all bins (see **Methods**).

The second phase in a DENR analysis is, optionally, to provide auxiliary information that may improve the accuracy of isoform abundance estimates. Any combination of three separate types of data can be provided: (1) the coordinates of predicted TSSs, (2) a list of inactive isoforms, and (3) a shape-profile correction. Separate predictions of TSSs are useful because they help to distinguish the start of one isoform (particularly one downstream from the start of a cluster) from the continuation of another isoform. The DENR package includes a pre-trained machine-learning classifier, implemented using TensorFlow, that can predict the locations of likely TSSs based on their characteristic patterns of bidirectional transcription and symmetric pause peaks (**Supplementary Figs. S2&S3;** see also ref. (24) for a similar approach). A separate specification of inactive isoforms is useful because it can direct the quantification algorithm to ignore a potentially large class of isoforms that may otherwise be misleading or confusing, based on auxiliary sources of data—including either experimental data, such as GRO-cap, PRO-cap, or RNA-seq, or computational predictions. Finally, the shape profile correction is a way of accommodating the typical “U”-shaped profile of nascent RNA sequencing reads along a gene-body, even after pause and termination peaks are excluded (**Fig. 1**). DENR provides a function to estimate the average profile from a designated subset of the data, and then to consider its shape when estimating the abundance of each isoform (see **Methods**).

Finally, DENR estimates the abundance of each isoform. Given the read counts per bin for each isoform cluster, DENR simply estimates a weight for each isoform by least squares, that is, by minimizing the squared difference between the expected density and the observed read count across all bins (see **Methods**). An option is also provided to perform this optimization in logarithmic space, i.e., by comparing the logarithm of the expected density and the logarithm of the read counts, corresponding to an assumption of a log-normal distribution for read counts (see **Discussion**).

### DENR accurately estimates RNA abundance at the gene and isoform levels

We evaluated DENR’s accuracy in quantifying RNA abundance at both the gene and isoform levels. Lacking an appropriate “gold-standard” in the form of real biological data, we chose to benchmark the software using simulated data. Because, to our knowledge, there is no available simulator for nascent RNA sequencing data that accommodates multiple isoforms per gene, we developed a new R package, called nascentRNASim, to provide a ground truth against which to compare DENR’s estimates (**Supplementary Fig. S4**). To make the simulated data as realistic as possible, nascentRNASim makes use of an empirical distribution of relative isoform abundances per gene obtained from RNA-seq data from GTEx (32). Given this distribution, the program then generates synthetic nascent RNA sequencing read counts for each isoform by resampling PRO-seq read counts from a manually curated set of archetypal transcripts (see **Methods**). The read counts from different isoforms are combined where they overlap. In this way, synthetic data is generated that closely resembles real data, without the need for restrictive modeling assumptions.

We first evaluated the impact of the various optional features by running the program with and without TSS prediction, shape-profile correction, log-transformation of read-counts, and with various numbers (0, 1, or 4) of masked bins at the 5′ and 3′ ends of each isoform. We ran DENR on 1500 simulated loci, measuring the Pearson’s correlation coefficient (*r*) of the estimated and “true” abundances at both the gene (**Supplementary Fig. S5**) and isoform (**Supplementary Fig. S6**) levels, and for dominant and longest isoforms as well as all isoforms together. We found, in general, that TSS prediction, the shape-profile correction, and the log tranformation did indeed improve performance significantly. The improvement was more substantial at the isoform level, where, together, these features increased *r* by as much as 10 – 15%, compared with an increase of ~3%. at the gene level. The effect of the masking strategy was more variable, but we found that masks of one bin at the 5′ end and four bins at the 3′ end performed well on average. Therefore, for all subsequent analyses (unless stated otherwise) on both simulated and real data, we used this masking strategy, and made use of TSS prediction, the shape-profile prediction, and the log-transformation.

With these options in place, we next compared DENR’s estimates for the same 1500 simulated loci with estimates obtained using a naive read-count-based (RCB) method commonly used in the field. For the RCB method, we simply estimated the abundance of a gene by the number of sequence reads that overlap any annotated isoform for that gene divided by the gene’s total length (see **Methods**). At the gene level, DENR’s estimates were highly concordant with true abundances (*r* = 0.97; **Fig. 2A**), substantially better than the RCB method (*r* = 0.85; **Fig. 2B**). Accordingly, DENR exhibited much smaller root-mean-square error (RMSE = 328.58) than the RCB method (RMSE = 642.19; **Fig. 2A&B**). DENR offered a particular improvement in cases where the dominant isoform corresponded to an internal TSS (**Supplementary Fig. S7A**), where the RCB method “over-normalized” using the length of whole gene and therefore underestimated abundance (**Supplementary Fig. S7B**; see **Supplementary Figs. S7C&D** for comparison). Notably, several genes having non-zero true abundances were estimated to have values of zero by DENR (**Fig. 2A**), apparently because of failures in TSS detection (see **Discussion**). The RCB method displayed the opposite tendency, estimating non-zero values for some genes having true values of zero (**Fig. 2B**). These cases were predominantly caused by overlap with or transcriptional run-on from other expressed genes.

**Fig. 2.**
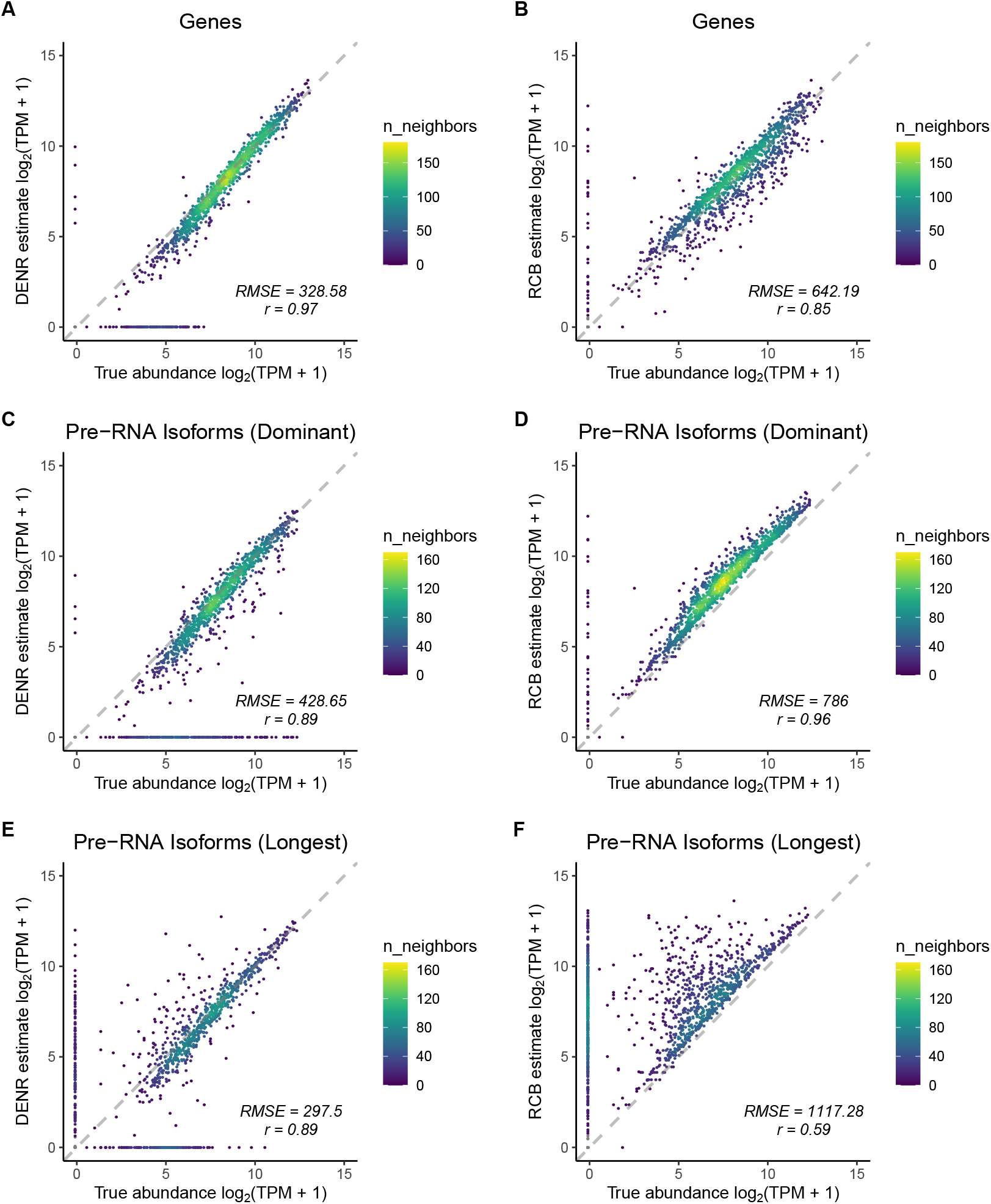
Comparison of DENR and the simple read-count-based (RCB) method for quantifying nascent RNA abundance. True (*x*-axis) vs. estimated (*y*-axis) abundance at the gene (**A & B**) and the isoform (**C–F**) levels, based on 1500 simulated loci. Data were simulated using nascentRNASim, which resamples real PRO-seq read counts and assumes a distribution of relative isoform abundances derived from real RNA-seq data. Results are shown for both the “ dominant” (most highly expressed) isoform (panels **C&D**) and the longest isoform (panels **E&F**). RMSE = root-mean-square error, *r* = Pearson’s correlation coefficient.

We also compared estimates from DENR and the RCB method with the true RNA abundances at the level of individual isoforms. We focused our evaluation on a single isoform per gene, selecting either the most abundant—or “dominant”—isoform, as determined by the true abundances; or the longest isoform, as determined by the annotations (see **Methods**). At the isoform level, DENR’s estimates of abundance were still well correlated with the true values (*r* = 0.89; **Fig. 2C**), although, not surprisingly, the concordance was somewhat reduced compared with the gene-level comparison (**Fig. 2A**). The estimates from the RCB method showed high correlation with true abundances (*r* = 0.96; **Fig. 2D**), but these estimates were systematically inflated, leading to substantially larger error (RMSE = 786) than that from DENR (RMSE = 428.65). This problem became more severe for the longest isoform, where DENR outperformed the RCB method substantially in terms of both correlation (*r* = 0.89 vs. 0.59) and RMSE (297.5 vs. 1117.28; **Fig. 2E&F**). These biases occur because the RCB method tends to misattribute sequence reads arising from other isoforms to the isoform in question. While other counting strategies could be devised, there is ultimately no good way to estimate isoform-specific abundance without simultaneously considering all candidate isoforms and all sequence reads (see **Discussion**).

### Application to real data for K562 and CD4^+^ T cells

Having demonstrated that DENR has good power to recover true gene and isoform abundances in simulated data, we next applied it to real data from K562 (25) and CD4^+^ T cells (33). We focused our analysis on 7732 and 7632 genes that displayed robust expression (ranking at top 75% of all expressed genes) in K562 and CD4^+^ cells, respectively. In K562 cells, we found that nearly half of these genes (3624 of 7732, or 46.9%) displayed evidence of expression at two or more isoforms (see **Methods**), indicating frequent use of alternative TSSs and/or TTSs (248 with alternative TSSs, 2213 with alternative TTSs, and 1163 with both). We observed a similar pattern in CD4^+^ cells, with 48.9% (3734 of 7632) of genes producing two or more pre-RNA isoforms. Moreover, we found that the dominant isoforms for 1178 (15.2%) and 1262 (16.5%) of genes, respectively, made use of an internal TSS, at least 1 kbp downstream from the 5′-most annotation.

To illustrate how DENR deconvolves the signal from PRO-seq data, we highlight two loci with multiple overlapping pre-RNA isoforms and evidence for internal TSS usage in K562 cells. The first example, at the gene *ST7*, is a relatively straightforward case (**Fig. 3**). This gene has 30 (mature-RNA) isoform annotations in Ensembl, which DENR merged into 19 distinct pre-RNA isoforms. However, the PRO-seq signal in the region suggests that only a subset of these isoforms are expressed, with clear signals beginning at a TSS near the 5′ end of the locus and at a second TSS about 60 kbp downstream. Indeed, DENR estimated nonzero abundance for only two isoforms, with the shorter one (G14406M1, corresponding to five Ensembl isoforms; see **Supplementary Table S1**) obtaining a higher weight than the longer one (G14406M6, corresponding to two Ensembl isoforms); the remaining 17 isoforms were assigned weights of zero. Notice that the TSSs of both isoforms are clearly marked by bidirectional transcription in the PRO-seq data, a signal used by DENR in picking them out.

**Fig. 3.**
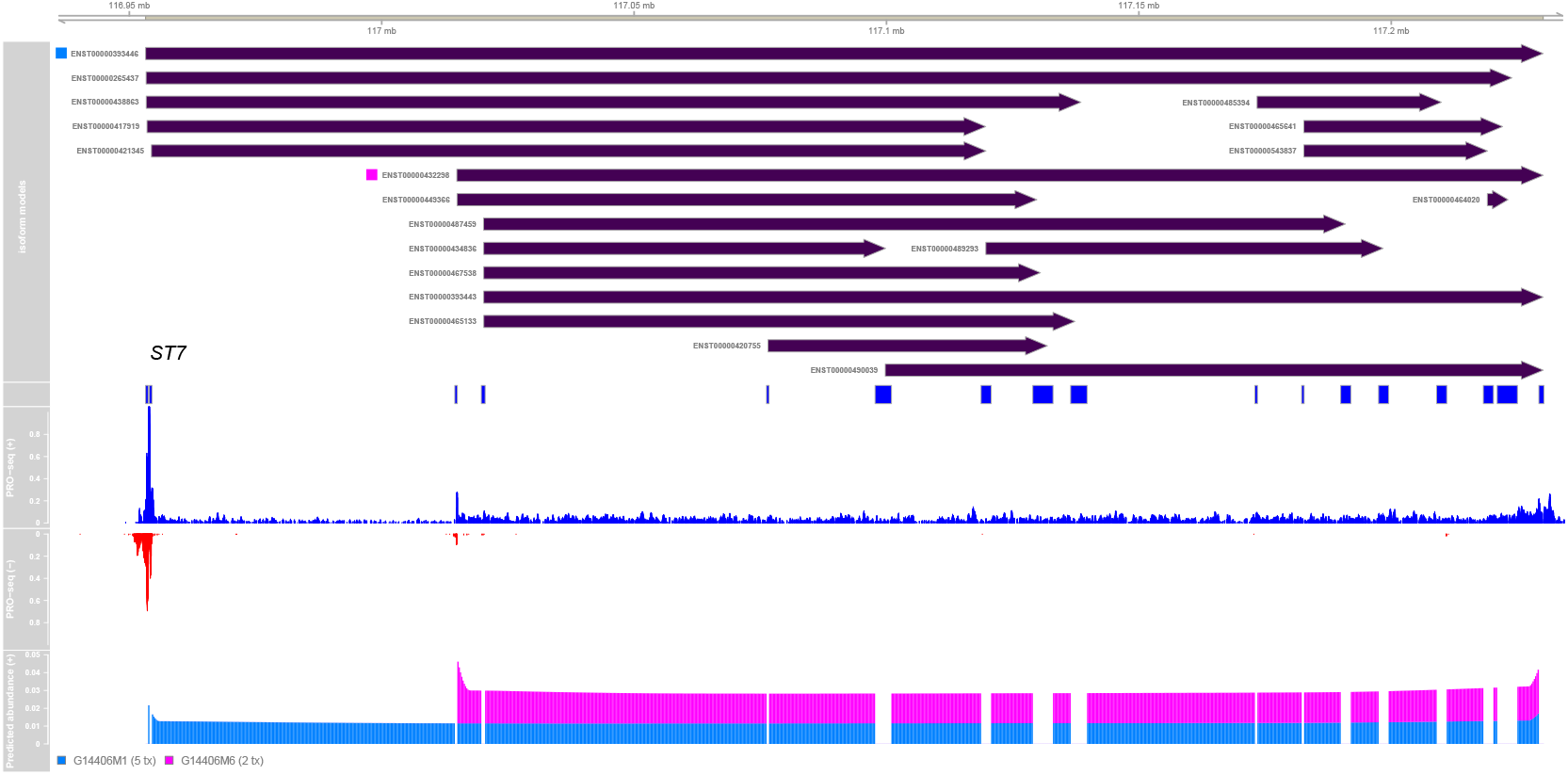
DENR abundance estimation for pre-RNA isoforms of *ST7* in K562 cells. The *ST7* (suppression of tumorigenicity 7; ENSG00000004866) gene has 30 isoform annotations in Ensembl, which DENR merges into 19 distinct pre-RNA isoform models (bars at top). Based on the observed PRO-seq data (*middle*, in blue and red), DENR estimates nonzero abundances for only two of these isoforms (marked in light blue and pink). The plot at *bottom* shows the expected relative contribution of each isoform model to the overall read counts per bin. Notice the effect of the shape-profile adjustment near the 5′ and 3′ ends. Notice also that the PRO-seq data reveals bidirectional transcription near the TSSs of both active isoforms; these signals are used by the machine-learning predictor to help identify sequence reads associated with these isoforms.

The second example is a more complex case in which three expressed genes (*SEC22C, SS18L2*, and *NKTR*) all overlap (**Fig. 4**). These genes all have multiple isoform annotations in Ensembl, some of which correspond to distinct pre-RNA isoforms after merging. In particular, *SEC22C* has 16 isoforms, which are merged into eight pre-RNA isoforms; *SS18L2* has three isoforms, which are merged into two; and *NKTR* has 19 isoforms, which are merged into ten. By again leveraging the signatures associated with TSSs, DENR identified two expressed isoforms of *SEC22C*, two expressed isoforms of *SS18L2*, and three expressed isoforms of *NKTR*. In each case, one isoform is clearly dominant, although in the case of *SS18L2*, both are expressed at non-negligible levels (**Supplementary Table S2**). Notice that the dominant isoforms for both *SEC22C* and *SS18L2* make use of internal TSSs. Notice also that DENR attributes both expressed isoforms of *SEC22C* and the minor expressed isoform of *SS18L2* to the same TSS, suggesting that stable transcripts are generated bidirectionally from this site. A second TSS contributes bidirectionally to the dominant isoform of *NKTR* and a minor isoform of *SEC22C*.

**Fig. 4.**
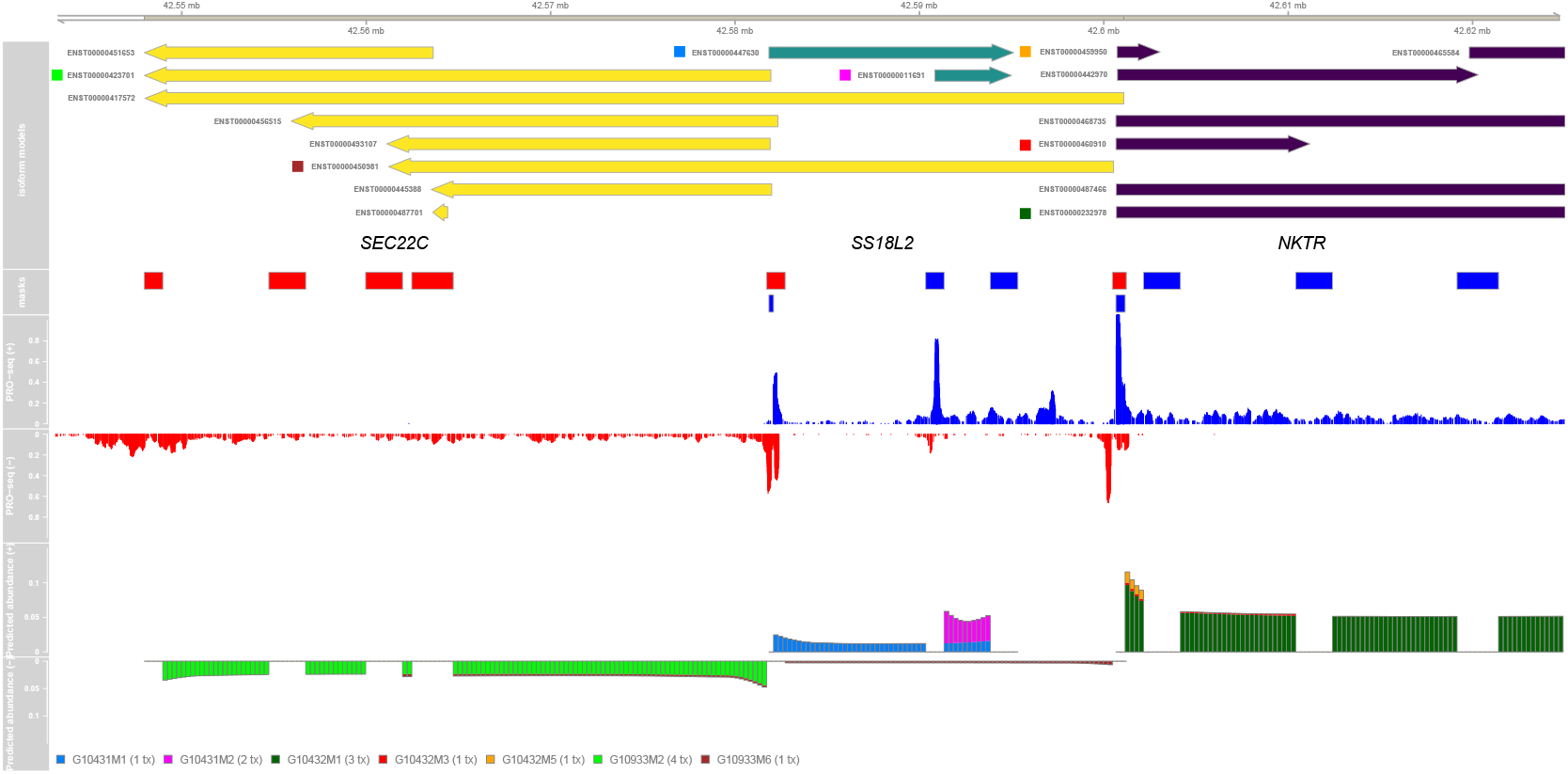
DENR abundance estimation for three overlapping genes on human chromosome 3. Isoform annotations are shown for *SEC22C* (yellow; ENSG00000093183), *SS18L2* (green; ENSG00000008324), and *NKTR* (purple; ENSG00000114857), together with the raw PRO-seq signal and DENR’s estimates of the expected contribution of each isoform model (with visualization conventions as in Fig. 3). Notice, again, the use of the shape-profile correction and the TSS predictions based on bidirectional transcription.

### Differences in dominant pre-RNA isoforms between CD4^+^ T cells and K562 cells

Given DENR’s ability to identify dominant pre-RNA isoforms, we wondered how frequently these isoforms might differ between cell types. We therefore compared the predictions of dominant isoforms from K562 cells to those from CD4^+^ T cells. Because the 3′ ends of pre-RNA transcription units can be difficult to pinpoint owing to transcriptional run-on, we focused on genes for which the dominant isoforms clearly used different TSSs in the two cell types, requiring a difference of at least 1 kbp in genomic coordinates (see **Methods**). In addition, we limited our analysis to 6757 genes showing robust expression (ranking in the top 75%) in both cell types. We found that 238 of these genes (~3.5%) had dominant isoforms that made use of different TSSs in K562 and CD4^+^ T cells. A gene ontology analysis showed that these genes were significantly enriched for annotations of alternative splicing (**Supplementary Fig. S8**), suggesting a correlation between alternative TSS usage and alternative splicing. One prominent example in this group is the gene encoding the transcription factor *RUNX1*, a master regulator of hematopoietic stem cell differentiation (**Fig. 5**), which has a much longer dominant isoform— resulting from a TSS about 160 kbp upstream—in CD4^+^ T cells as compared with K562 cells. This gene is known to make use of alternative TSSs in a temporal and tissue-specific manner (34–36). Additional examples are shown in **Supplementary Figures S9&S10.**

**Fig. 5.**
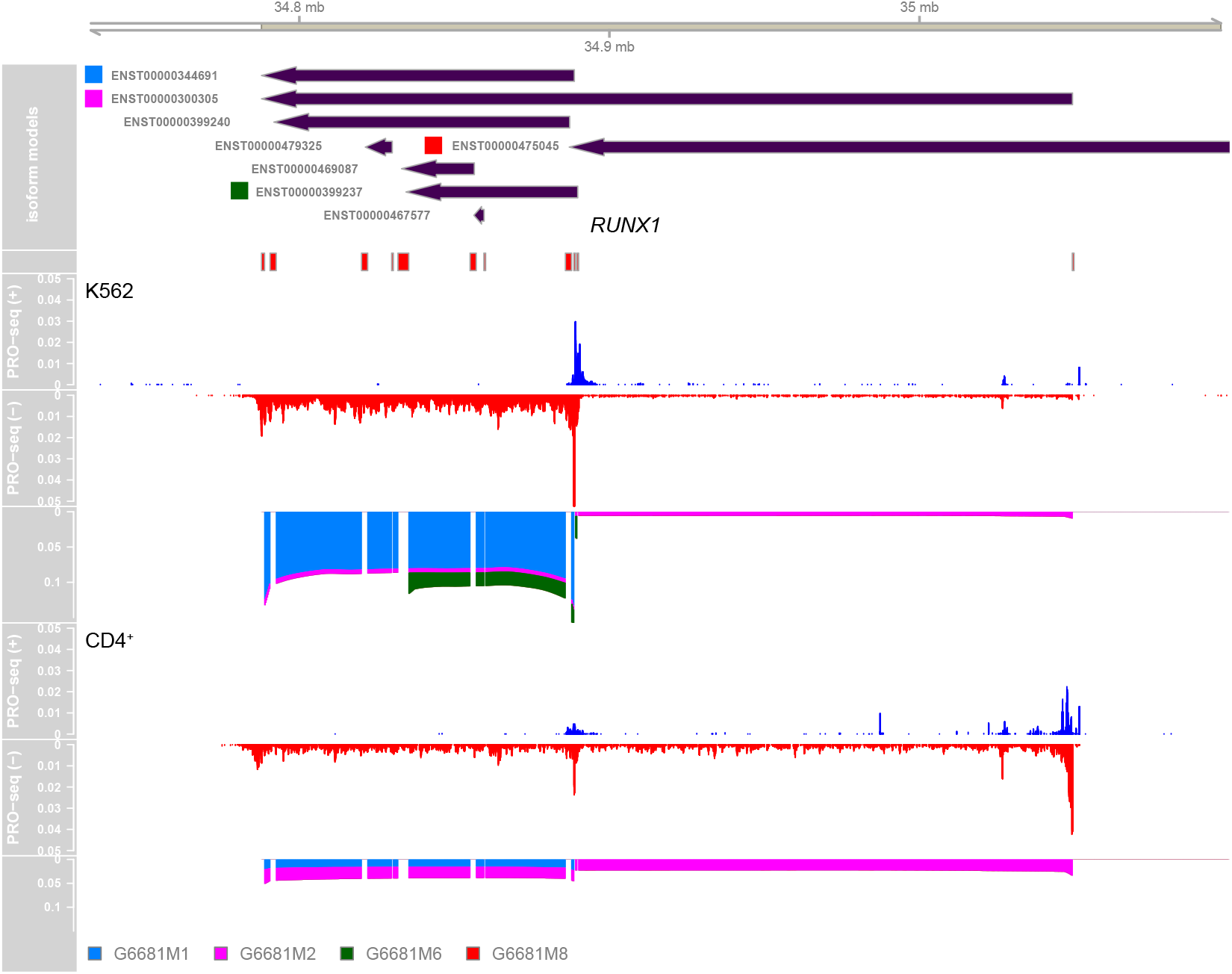
Cell-type specific TSS usage for *RUNX1*. Of several annotated pre-RNA isoforms for the gene encoding the transcription factor RUNX1 on human chromosome 21 (shown on the negative strand at *top*), DENR finds two isoforms to be dominant: a ~100-kb isoform (G6681M1; shown in blue), and an isoform that is more than twice as long and begins ~160 kb upstream (G6681M2; shown in pink). The shorter isoform is clearly dominant in K562 cells (*middle*), whereas both are expressed at non-negligible levels in CD4^+^ T cells, with the longer one being slightly dominant (*bottom*). RUNX1 is essential for normal hematopoietic development and its dysregulation is associated with hematological malignancies (34). It is well known to make use of alternative promoters (35,36)

### Relative contributions of transcriptional and post– transcriptional processes to isoform diversity

We were interested in making use of DENR to assess overall levels of isoform diversity genome-wide. Furthermore, we wondered if a parallel analysis of RNA-seq data would enable an informative comparison of the relative contributions to isoform diversity at the pre-RNA and mature RNA levels. Toward this end, we generated high-quality matched PRO-seq and RNA-seq data sets (both with paired-end reads; see **Methods**) for two similar but distinct human cell types, CD4^+^ T cells and CD14^+^ monocytes. We used DENR to quantify isoform diversity at the pre-RNA level and StringTie (37) to quantify isoform diversity at the mature RNA level in each cell type. Isoforms not detected in RNA-seq were also used to indicate non-active isoforms in DENR instead of using TSS prediction. Finally, we focused our analysis on a set of 10,650 genes that were expressed in both cell types, with good representation in both the PRO-seq and RNA-seq data sets (see **Methods**).

To quantify isoform diversity at the pre-RNA and mature RNA levels, we made use of the information-theoretic measure of Shannon entropy. We observed that, given pre-RNA isoform abundance relative frequencies *X* (estimated from PRO-seq data using DENR) and mature RNA isoform abundance relative frequencies *Y* (estimated from RNA-seq data using StringTie), the joint entropy *H*(*X,Y*) can be decomposed into a component arising from primary transcription, *H*(*X*), and a conditional-entropy component arising from post-transcriptional processes, *H*(*Y*|*X*); that is, *H*(*X,Y*) =*H*(*X*) +*H*(*Y*|*X*) (see **Methods**). Thus, we can estimate *H*(*X*) across any set of expressed genes using DENR, estimate *H*(*X*,*Y*) for the same set of genes using StringTie, and then estimate the post-transcriptional entropy, *H*(*Y*|*X*) by their difference. We can further estimate the fractional contribution of transcription to the final isoform entropy as *H*(*X*)/*H*(*X,Y*).

When applying these methods to the CD4^+^ T cell and CD14^+^monocyte data sets individually, we observed reasonably good concordance, with estimates of *H*(*X,Y*) = 0.94−1.01 bits/gene in total entropy, of which 63–64% comes from transcriptional entropy (*H*(*X*)) and the remaining 36–37% derives from post-transcriptional processes (**Fig. 6A&B**). When we pooled data from the two cell types together (“both”), *H*(*X,Y*) increased by about 10%, indicating higher levels of isoform diversity across cell types than within them. Interestingly, however, the fractional contribution from primary transcription, *H*(*X*)/*H*(*X,Y*), also increased substantially, from ~0.64 to ~0.72, suggesting that transcriptional processes make a disproportional contribution to the isoform diversity across cell types, which is more likely than diversity within each cell type to be associated with true functional differences (see **Discussion**).

**Fig. 6.**
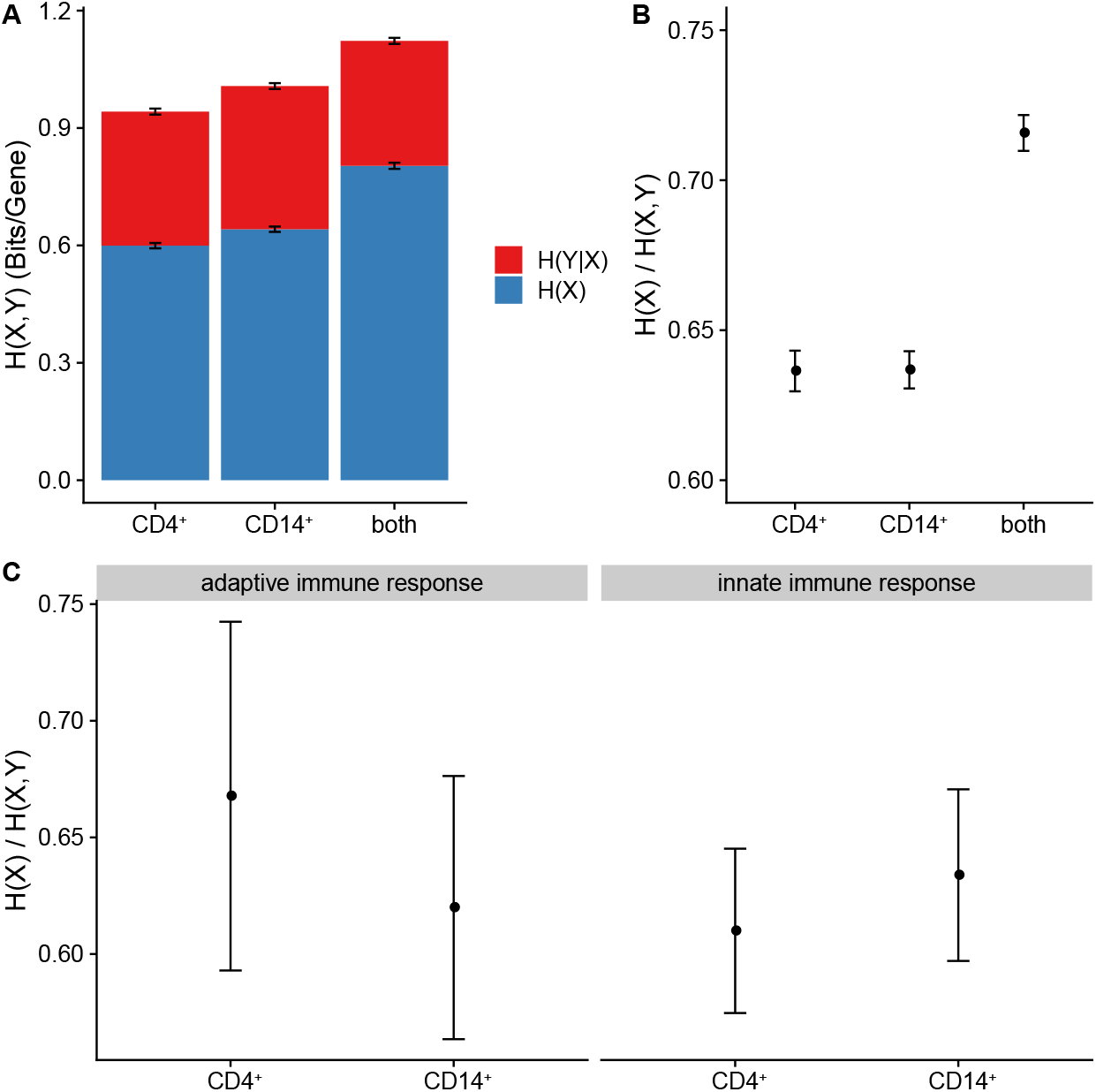
Decomposition of Shannon entropy of isoform diversity into contributions from primary transcription and post-transcriptional processing. **(A)** Entropy per gene of mature RNA isoforms (*H*(*X,Y*)) is partitioned into a component from primary transcription (*H*(*X*)) and a component from post-transcriptional processing, including splicing (*H*(*Y|X*)). **(B)** Fractional contribution from primary transcription, *H*(*X*)/*H*(*X,Y*). Results are for 10,650 genes expressed in both CD4^+^ T cells and CD14^+^ monocytes. “Both” indicates results when both data sets are pooled. **(C)** Fractional contribution from primary transcription, as in **(B)**, but for the subsets of genes associated with the Gene Ontology terms “adaptive immune response” (GO:0002250; *n* = 116) and “innate immune response” (GO:0045087; *n* = 287). Error bars represent the standard deviation of the mean as estimated by bootstrap resampling (*n* = 100).

A primary difference between these cell types is that CD4^+^T cells play an important role in the adaptive immune system whereas CD14^+^ monocytes are part of the innate immune system. Therefore, we extracted 116 and 287 genes associated with the Gene Ontology (GO) terms “adaptive immune response” and “innate immune response,” respectively, and calculated *H*(*X*)/*H*(*X,Y*) separately for each of these these subsets of genes. Interestingly, we found that this fraction was somewhat elevated in adaptive-immunity-related genes in CD4^+^ T cells, and slightly elevated in innate-immunity-related genes in CD14^+^ monocytes (**Fig. 6C**), suggesting that primary transcription may disproportionally contribute to isoform diversity in the genes most relevant to the specific immune-related functions of each cell type.

## Discussion

In this article, we have introduced Deconvolution of Expression for Nascent RNA-sequencing data (DENR), the first fully vetted computational method—to our knowledge—to address the abundance estimation problem at the level of pre-RNA isoforms, based on nascent RNA sequencing data. At its core, DENR is simply a regression-like method for estimating a weight for each element in a set of predefined candidate isoforms, by minimizing the sum-of-squares difference between expected and observed read counts. This baseline model is augmented by various refinements, including machine-learning predictions of transcription start sites, a shape-profile correction for read counts, and masking of read counts near isoform TSSs and TTSs. We have shown that DENR performs well on simulated and real data, and can be used for a variety of downstream applications.

In direct comparisons with simple read count-based (RCB) methods like those used in most current applications, we find that DENR does indeed offer a substantial performance improvement. The improvement is most pronounced at the isoform level, where the RCB methods inevitably misattribute many reads to the wrong isoform. Interestingly, however, DENR also improves substantially on gene-level estimates of abundance. The main reason for this improvement has to do with the normalization for gene length. The gene-level RCB method has no good way to identify which bases in the DNA template are transcribed, and must conservatively assume transcription occurs across the union of all annotated isoforms. As a result, it frequently “over-normalizes” and underestimates abundance. DENR, by contrast, simultaneously models all isoforms and explains the full set of read counts at a locus as a mixture of isoforms. The limitations we observed with alternative RCB methods highlight the difficulty of accurately estimating abundance without a model that assigns reads to isoforms in zero-sum fashion. Because most reads can potentially arise from multiple alternative isoforms, any naive counting method will tend to either over- or under-estimate abundance. These errors in abundance estimation, in turn, can result in biases in many downstream applications, such as elongation-rate or RNA-half-life estimation.

In analyses of real data, we found that many genes (nearly half of robustly expressed genes in K562 and CD4^+^ T cells) display evidence of expression at multiple distinct pre-RNA isoforms. Moreover, we found that the dominant isoform fairly commonly (in ~15% of cases) makes use of a TSS that is substantially downstream of the 5′-most annotation. These cases are particularly likely to be mischaracterized by standard methods for quantifying pre-RNA expression. We have highlighted specific examples showing how DENR can effectively deconvolve the read-count contributions of multiple overlapping isoforms, including a complex case involving multiple overlapping genes (**Fig. 4**). In addition, in a comparison of K562 and CD4^+^ T cells, we identified more than two hundred genes that use different dominant isoforms in these two cell types, including prominent examples such as *RUNX1*.

One interesting consequence of having the ability—as we now do—to characterize the distribution of isoform abundances at both the pre- and mature-RNA levels is that it potentially allows for a decomposition of the contributions to isoform diversity from primary transcription and post-transcriptional processes. In a final analysis, we attempted to quantify these relative contributions using a simple information theoretic calculation, by partitioning the Shannon entropy in mature-RNA isoform diversity (as estimated from RNA-seq data using StringTie) into a component estimated at the pre-RNA level (by applying DENR to PRO-seq data) and the remainder, which we argue can be interpreted as the conditional entropy introduced at the post-transcriptional level. Our observations are qualitatively similar to a number of previous studies reporting observations of widespread, regulated alternative TSS usage, often in a tissue-specific manner (38–40), some of which have argued for a primary role of transcription relative to splicing (41,42). However, while the post-transcriptional entropy that we measure presumably derives primarily from splicing, it is worth noting that it could also be influenced by post-transcriptional up- or down-regulation of particular isoforms, for example, through miRNA- or RBP-mediated decay. In some cases, post-transcriptional processes could even reduce entropy generated at the pre-RNA level, for example, by sharply down-regulating particular pre-RNA isoforms relative to others. Importantly, this type of generation or reduction in entropy can only be detected if pre-RNA isoform diversity is independently characterized by a method like the one introduced here, rather than indirectly assessed from RNA-seq (or CAGE) data. For this reason, we believe our analysis is complementary to previous analyses of alternative promoters and TSSs.

There are a number of potential avenues for improvement of our current implementation of DENR. First, the method assumes a sum-of-squares loss function, which is equivalent to maximum likelihood estimation under a Gaussian (or log normal, if optimized in log space) generating distribution for read counts, with the counts for each bin assumed to be independent and identically distributed. Real nascent RNA sequencing read counts, however, tends to be not only overdispersed but nonuniform along the genome, with fairly pronounced spikes separated by intervals of reduced signal. The method could be extended to allow for maximum-likelihood estimation under an arbitrary generating distribution for read counts, by making use of a general probabilistic model for nascent RNA sequencing data that we have recently proposed (43). This model could potentially accommodate autocorrelated read counts along the genome sequence, although in this case, optimizing the mixture coefficients would become more complex and computationally expensive. Another advantage of this framework is that it would naturally accommodate a richer and more general model for changes in polymerase density along the gene body, beyond the simple shape-profile correction introduced here. As a result, it might require a less heavy-handed masking strategy, by providing a better description for read counts near TSSs and TTSs. More work will be needed to determine if these generalizations are sufficiently advantageous to justify their complexity and computational costs.

A second limitation is that DENR effectively uses a “hard prior” for candidate isoforms, either treating them as equally likely *a priori* or completely excluding them (i.e., assigning a prior probability of zero) based on the absence of a TSS prediction or other evidence of inactivity. A natural generalization would be to accept an arbitrary prior probability for each candidate isoform. These weights could potentially be determined based on a variety of relevant covariates, including not only TSS predictions but also, say, chromatin accessibility, chromatin contact, histone modification, or RNA-seq data from a relevant cell type. The model would then combine the prior probabilities with the data likelihood to enable full Bayesian estimation of isoform abundances. A related extension would be to consider not only annotated isoforms but also ones suggested by the nascent RNA sequencing data but not annotated. Such candidates could potentially be identified using a separate method (e.g., (44)) and given lower prior weights than annotated isoforms; if they had sufficient support in the data, they might still obtain high posterior probabilities.

Finally, the current inference method does not make use of a sparsity penalty to encourage the observed data to be explained using as few isoforms as possible. In initial experiments, we did not find that such penalties made a noticeable difference in our prediction performance, and in general, we do not observe a proliferation of isoforms with small weights. However, we do occasionally find that DENR gives high weights to short transcripts that happen to coincide with spikes in the data or pause peaks, apparently owing to a failure to account for spikes in the read-count data, as well as inadequacies in the shape-profile correction when applied to short isoforms. It is possible that a sparsity penalty—perhaps combined with the use of a richer model for read counts—would help to eliminate some of these apparently spurious predictions.

Despite these limitations, we have shown that DENR is generally an effective tool for quantifying pre-RNA abundance at both the gene and isoform levels, with many possible down-stream applications. We expect this method to be increasingly useful to the community as nascent RNA-sequencing data grows more abundant and is used for a wider variety of downstream applications.

## Methods

### Estimating isoform abundance

DENR estimates the abundance of each isoform by non-negative least-squares optimization, separately at each cluster. For a given cluster of *n* isoforms spanning *m* genomic bins, let *β* = (*β*_1_,…, *β_n_*)′ be a column vector representing the coefficients (weights) assigned to the isoforms, let **Y** = (*y*_1_,…,*y_m_*)′ be a column vector representing the read-counts in the bins, and let **X** be an *n* × *m* design matrix such that *x*_*i,j*_ = 1 if isoform *i* spans bin *j* and *x_i,j_*, = 0 otherwise. DENR estimates *β* such that,

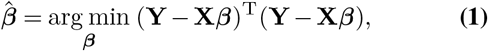

subject to the constraint that *β_i_* ≥ 0 for all *i* ∈ {1,…,*n*}. If the option to apply a log-transformation is selected, then the transformation is applied to both the elements of **Y** and those of **X***β*, and the optimization otherwise proceeds in the same manner. In either case, DENR optimizes the objective function numerically using the BFGS algorithm with a boundary of zero for the *β_i_* values. Notice that, when the shape-profile correction is applied, the non-zero values in the design matrix **X** are adjusted upward and downward from 1 (see below).

After obtaining estimates for all isoform abundances *β_i_*, we normalize them by the total library depth to facilitate comparisons between samples. Isoform-level abundances are then converted to gene-level abundances by summing over all isoforms associated with each gene.

### Machine-learning predictor for active TSSs

To distinguish active and inactive TSSs based on patterns of bidirectional transcription in nascent RNA-sequencing data, we implemented a convolutional neural network (CNN) classifier using the Keras interface to TensorFlow (45). We trained the CNN on previously published PRO-seq data from K562 cells (25), using matched GRO-cap data (23) to identify positive and negative examples. Specifically, we conservatively defined candidate TSSs as ‘active’ if they were overlapped with GRO-cap peaks from a HMM-based predictor described in ref. (23), and TSS with max GRO-cap signal was selected per peak; in addition, we defined candidates as ‘inactive’ if TSSs were not overlapped with any peaks by the HMM-based predictor and no active TSSs (distance > 100*bp*) and raw GRO-cap signal (distance > 25*bp*) nearby (**Supplementary Fig. S2**). The CNN was composed of two 1-D convolutional layers, the first with a ReLU activation function and both with max-pooling, followed by drop-out, a densely connected layer, and a single sigmoid output (**Supplementary Fig. S3**). It was applied to feature vectors corresponding to strand-specific read counts in 21 bins of width 51 bp, centered on the positive and negative examples; the 42 raw read-counts for each example were transformed to *z*-scores for scale-independence. The CNN was trained using the Adam optimizer (46) with early stopping.

When the optional TSS-calling feature is in use, only isoforms corresponding to predicted ‘active’ TSSs are allowed to have non-zero weights. However, because the TSS predictor inevitably misses some active TSSs that exhibit unusual patterns of aligned reads, DENR makes use of a heuristic method to identify and reconsider regions of “unexplained” high-density polymerase. Specifically, an upstream polymerase ratio (UPR) statistic is calculated by taking the ratio of the read-count density immediately upstream of the gene and inside the gene body. If the UPR of an isoform is ≥10, and there are no other active isoforms within 5 kbp upstream or 6 kbp downstream of its TSS, then the isoform is eligible to be assigned a non-zero weight.

### Shape-profile correction

The shape-profile correction is empirically derived from a reference set of isoforms. Briefly, starting with the full set of annotations provided by the user, DENR identifies a subset of isoforms that, according to various heuristics, appear to be sufficiently long, robustly expressed, and the sole source of sequencing reads in their genomic regions. DENR then tiles each representative isoform with bins of the user-specified size (default 250 bp), and maps those bins to a canonical [0, 1] interval. This mapping is intended to fix the scales of the promoter-proximal and termination regions, and allow the remaining gene-body to be compressed or expanded as needed. Specifically, the first 15 kbp of each isoform is mapped (proportionally) to the interval [0, 0.2], the last 5 kbp is mapped to [0.8, 1], and the remaining portion is mapped to the (0.2, 0.8) interval. Finally, the canonical shape-profile is obtained by averaging the relative read-count densities of the entire [0,1]-rescaled reference set of isoforms, using a loess fit for smoothing, and scaling the density such that the median value across the entire interval is one. This shape-profile is then used to adjust the design matrix **X** (see above) by replacing each value of 1 with the relative density at the corresponding location in the canonical shape profile. The isoform weights are then estimated by least-squares, as usual. In the case of isoforms of length *l* ≤ 20 kbp, the first 0.75*l* and last 0.25*l* base-pairs are proportionally mapped to the [0, 0.2] and [0.8, 1.0] intervals, respectively, in the canonical shape-profile, and the interval (0.2, 0.8) is ignored.

### Simulation of nascent RNA sequencing data

Our nonparametric simulator for nascent RNA sequencing data, called nascentRNASim, makes use of a template set of isoform annotations and a designated collection of well-defined isoform “archetypes” and corresponding read counts. Given these inputs, we simulate a synthetic data set in five steps. First, we group the isoform annotations into non-overlapping strand-specific clusters, as in a DENR analysis. Second, we sample randomly (with resampling) from this set of clusters, and similarly, from the set of inter-cluster distances. Third, within each sampled cluster, we substitute for each isoform the archetype that is closest to it in genomic length, keeping the TSS at its original position relative to the beginning of the cluster. Fourth, we sample a new isoform abundance for each synthetic isoform from a distribution fitted by kernel density estimation to isoform abundance estimates from GTEx for skeletal muscle (32). Finally, we obtain a new read count for each bin along the isoform by resampling from the original value in proportion to the simulated abundance estimate. In this way, we sample a full synthetic data set, consisting of realistic clusters, each with a realistic distribution of isoforms and realistic patterns of read counts, but with a known abundance for each isoform.

In this work, we used the PRO-seq data set from ref. (25) as our source data set, together with isoforms from Ensembl. We selected a set of 62 archetypes manually, looking for isoforms with a range of lengths that exhibited relatively high read depth, appeared to be solely responsible for the local PRO-seq signal (i.e., they did not overlap other active isoforms and were at least ~5 kbp from other active genes), and showed a PRO-seq signal that approximately coincided with the annotated TSS and TTS, dropping to background levels nearby. We also considered GRO-cap data from ref. (23) in identifying TSSs. Notice that the design of the simulator ensures that every synthetic isoform has the same length and approximate read-count pattern as one of the 62 archetypes, but multiple isoforms may overlap (with additive contributions to read counts) in the synthetic data.

### Calculation and decomposition of Shannon entropy

Let *X_i_* be a random variable representing the possible pre-RNA isoforms of gene *i*, and assume the probability density function for *X_i_* is proportional to DENR-based estimates of isoform abundance. That is, 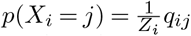, where *q_ij_* is the estimated abundance of the *j*th isoform of gene *i* and 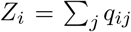. We calculate the Shannon entropy of *X_i_* as 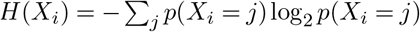, and we calculate the total entropy of a set of genes *S* as 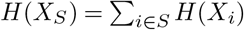, assuming independence of genes.

Similarly, let *Y_i_* represent the possible mature RNA isoforms of gene *i*, with 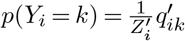, where 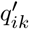 is the StringTie-estimated abundance of the *k*th isoform of gene *i* and 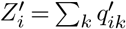. Then 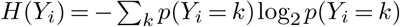, and, for a set of genes *S*, 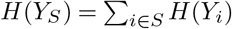.

To decompose entropy into components from *H*(*X*) (primary transcription) and *H*(*Y*|*X*) (post-transcriptional processes), we consider the joint entropy of *X* and *Y*, *H*(*X,Y*), and make use of the chain rule for joint and conditional entropy, *H*(*Y*|*X*) = *H*(*X,Y*) − *H*(*X*), interpreting *H*(*Y*|*X*) as the additional entropy contributed to the distribution of pre-RNA isoforms by post-transcriptional processes. Furthermore, because in this case, each mature RNA isoform corresponds to a single pre-RNA isoform, *H*(*X,Y*) is the same as *H*(*Y*). Specifically, for each *i*,

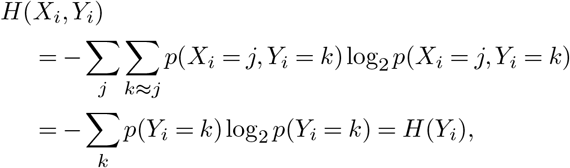

where *k* ≈ *j* indicates that mature RNA isoform *k* is compatible (in TSS and TTS) with pre-RNA isoform *j*.

### Human blood CD14^+^ monocytes and CD4^+^ T-cells isolation

Roughly 80 ml of human blood was drawn to and kept in spray-coated EDTA tubes (BD Vacutainer #366643) at 4°C. The following day, blood was diluted 1:1 in PBS and loaded on an equal volume of Ficoll-Paque (Fisher Scientific #45-001-750). Peripheral blood mononuclear cells (PBMCs) were then isolated by centrifugation for 20 min at 750xg. All layers, excluding the lower erythrocytes-containing layer, were moved to a new 50ml tube, and washed twice in PBS. PBMCs were then treated with 2 ml of erythrocytes lysis buffer (Lonza Walkersville inc. #120-02-070) for 1 min, flooded with 5 ml of RPMI supplemented with 10% FBS, and centrifuged for 10 min at 4°C and 1500 RPM on a Megafuge 40R Refrigerated Centrifuge (Thermo Scientific #75004518). CD14^+^ monocytes were isolated from PBMCs by a magnetic cell separation system (MACS) using anti-human CD14 antibody attached to microbeads (Miltenyi Biotec #130050201) on an LS column (Miltenyi Biotec #130-042-401) following the manufacturer’s protocol, while maintaining the flowthrough of PBMCs without CD14^+^ monocytes. CD4^+^ T cells were then isolated from CD14^+^ monocyte-free PBMCs using anti-human CD4 antibody attached to microbeads (Miltenyi Biotec #130045101) on a new LS column, following the same protocol. Finally, CD14^+^ monocytes and CD4^+^ T-cells were incubated for 1h of recovery in RPMI supplemented with 10% FBS before any downstream applications.

### RNA-seq library preparation

Cells were flooded with 1 ml per 5 × 10^6^ cells of TRI reagent (Molecular Research Center #TR118) and 0.2 ml of chloroform was added per 1 ml of TRI reagent followed by a vigorous vortexing for 20 sec. Cells were then incubated in the TRI reagent and chloroform solution for 2 minutes, followed by a 12,000xg centrifugation at 4°C for 15 min. The resulting aqueous phase was transferred to a new 1.5 ml tube and 0.5 ml of isopropyl alcohol was added for each 1 ml of TRI reagent initially used, and the solution was incubated in room temperature for 10 min, followed by a 10 min centrifugation in 12,000xg at 4°C. RNA was then washed twice with 75% ethyl alcohol, air-dried for 5 min with an open lid on ice and dissolved in DEPC-treated water. Poly-A enriched RNA-seq libraries were then prepared with up to 1 μg of isolated RNA, using the NEBNext^®^ Ultra^™^ II Directional RNA Library Prep Kit for Illumina®(New England Biotech #E7760) with the NEBNext Poly(A) mRNA Magnetic Isolation Module (New England Biotech #E7490), following the manufacturer’s protocol.

### Nuclei isolation and PRO-seq library preparation

1 × 10^6^ cells were centrifuged at 4°C for 5 min at 1000xg and washed twice with 1 ml of PBS. Cells were then re-suspended in 150 μl of wash buffer (10mM Tris-Cl pH 8.0, 300mM sucrose, 10mM NaCl, 2mM MgAc_2_, 2.5 μM DTT, 1X protease inhibitor cocktail (Thermo Scientific, #A32965)) supplemented with 0.6U of SUPERase In RNase Inhibitor (Thermo Fisher Scientific, #AM2696). 150 μl of 2X lysis buffer (10mM Tris-Cl pH 8.0, 300mM sucrose, 10mM NaCl, 2mM MgAc_2_, 6mM CaCl_2_, 0.2% NP-40) were added to the solution and samples were gently pipetted up and down 10 times, to facilitate nuclei release. Released nuclei were then centrifuged at 4°C for 5 min at 1000xg, washed with 1 ml of a 1:1 ratio solution of wash buffer and 2X lysis buffer and re-suspended in 50 μl of storage buffer (50mM Tris-CL pH 8.3, 40% glycerol, 5mM MgCl_2_, 0.1 mM EDTA, 2.5 μM DTT, 1X protease inhibitor cocktail) supplemented with 0.2U of SUPERase In RNase Inhibitor. PRO-seq run-on and library preparation was completed following a recently updated protocol (47). Briefly, nuclei were run-on by incubating at 37°C for 5 min in run-on buffer (10 mM Tris-Cl pH 8.0, 5 mM MgCl_2_, 1 mM DTT, 300 mM KCl, 40 μM Biotin-11-CTP, 40 μM Biotin-11-UTP, 40 μM Biotin-11-ATP, 40 μM Biotin-11-GTP, 1% (w/v) Sarkosyl in DEPC H_2_O). The run-on reaction was stopped by adding Trizol LS (Life Technologies, #10296-010). RNA was pelleted with the addition of GlycoBlue (Ambion, #AM9515) to visualize the pellet, re-suspended in diethylpyrocarbonate (DEPC)-treated water and heat denatured for 40 sec. RNA was digested using 0.2N NaOH on ice for 6 min, which yields RNA lengths ranging from ~20-500 bases. The 3′ adapter (sequence: AGATCGGAAGAGCACACGTCTGAACTC) was ligated using T4 RNA Ligase 1 (NEB, M0204L) and purified nascent RNA using streptavidin beads (NEB, S1421S). Next the nascent RNA was decapped using RppH (NEB, M0356S), the 5′ end was phosphorylated using T4 polynucleotide kinase (NEB, M0201L), and the 5′ adapter (sequence: GTTCAGAGTTC-TACAGTCCGACGATC) was ligated. RNA was removed from the beads and a reverse transcription was performed using Superscript IV Reverse Transcriptase (Life Technologies) and amplified using Q5 High-Fidelity DNA Polymerase (NEB, M0491L). PRO-seq libraries were sequenced using the NextSeq500 high-throughput sequencing system (Illumina) at the Cornell University Biotechnology Resource Center.

### Applying DENR to synthetic data

To benchmark DENR’s performance, nascentRNASim was first used to simulate PRO-seq read-counts for 1500 genes. To thoroughly examine the effects of optional features on performance, all combinations of optional features, i.e., with and without TSS prediction, shape-profile correction, log-transformation of read-counts, and with various numbers (0, 1, or 4) of masked bins at both the 5′ and 3′ end of each isoform, were tested on the synthetic data, resulting in a total of 72 test schemes (2^3^ × 3^2^; **Supplementary Figs. S5&S6**). The scheme with TSS prediction, shape-profile correction, log-transformation of read-counts, masking of one bin around the TSS and four bins around the TTS performed well at both the gene and isoform levels. Therefore, this combination was used for all subsequent analyses in synthetic and real data with one exception: in the entropy analysis, we used RNA-seq data indicate non-active isoforms instead of relying on TSS prediction. The gene-level comparison was performed on the whole set of genes, and on two complementary subsets: one for which active isoforms predominately used an internal TSS, and one for which they used the 5′-most TSS for transcription. Genes were defined as using an internal TSSs if their dominant isoforms were transcribed from a TSS at least 1 kbp downstream from the 5′-most TSS annotation; otherwise they were defined as using the 5′-most TSS (**Supplementary Fig. S7)**. At the isoform level, we compared the performance of DENR and the RCB method for both dominant isoforms determined by true abundances in simulation, and longest isoforms determined by the annotations. To make the estimates more comparable, we masked 250 bp downstream from TSS and 1000 bp upstream from the TTS when counting reads for the RCB method.

For the RCB method, the abundance of a gene or isoform *i* is estimated as follows:

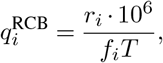

where *r_i_* is number of reads mapped to the genomic region in question (corresponding either to an isoform or the union of isoforms associated with a gene), *f_i_* is the length of that region, and,

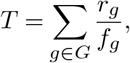

where *G* is the set of all genes in the simulation, *r_g_* is number of reads mapped to a gene region, and *f_g_* is the length of that gene. Notice that 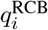 has units of transcripts per million (TPM) (48).

### Applying DENR to real data

To prepare bigWig files as input for DENR, published K562 (25) and CD4^+^ T cell (33) PRO-seq libraries were first processed by the PROseq2.0 pipeline using single-end mode (49). The human genome assembly (GRCh38.p13) and isoform annotations were downloaded from Ensembl (release 99) (50). Annotations of protein-coding genes from the autosomes and X chromosome were used, excluding genes that overlapped on the same strand. To identify genes producing two or more pre-RNA isoforms with high confidence, only genes with robust expressions (i.e., ranking at top 75% of all expressed genes) in K562 (*n* = 7732) and CD4^+^ T cells (*n* = 7632) were retained for analysis. To survey predominant usage of internal TSSs for transcription, genes with dominant pre-RNA isoforms transcribed from internal TSSs 1 kbp downstream from the 5′ most TSSs were identified and visualized using Gviz (51).

To investigate the differences in dominant isoforms between K562 and CD4^+^ T cells, mature RNA isoform annotations were first grouped together if the distances between their annotated TSSs were <1 kbp. The longest isoform in each group was selected as the representative and used for estimating abundance. Inactive TSSs were predicted separately in K562 and CD4^+^ T cells then intersected, to ensure that the same set of inactive isoforms was used across cell types. To identify genes with different dominant isoform between cell types, 6757 genes exhibiting robust expression (i.e., ranking in the top 75% in both cell types) were analyzed. We focused on cases in which the dominant isoforms differed in the two cell types. Gene Ontology analysis was performed using the online tool DAVID (52).

For calculation of Shannon entropy in the newly generated, matched PRO-seq and RNA-seq data for CD4^+^ T cells and CD14^+^ monocytes, we estimated isoform abundances at the pre-RNA and mature RNA levels. For mature RNA abundances in RNA-seq, adapters (sequence: AGATCGGAA-GAGC) in raw reads were first removed using Cutadapt (v2.10) (53). Clean reads were then aligned to the human genome using HISAT2 (v2.2.1) (54). Mature RNA isoform abundances were quantified using StringTie (v2.1.4) (37). Human genome (GRCh38.p13) and GTF files were downloaded from Ensembl (release 99) (50). For pre-RNA isoform abundances, PRO-seq libraries were first processed by the PROseq2.0 pipeline using paired-end mode (49), then DENR was used to quantify abundances. The shape-profile correction, log-transformation of read-counts, and masking of one around the TSS and four bins around the TTS were used, as in other analyses, but that inactive isoforms were identified as those not detected in the RNA-seq data rather than by using DENR’s TSS prediction feature. 10,650 genes with abundance estimates > 0 in CD4^+^ T cell and CD14^+^ monocyte samples were used for the entropy calculation.

### Software availability

DENR and nascentRNASim are freely available at https://github.com/CshlSiepelLab/DENR (version v1.0.0) and https://github.com/CshlSiepelLab/nascentRNASim (version v0.3.0). Usage of DENR is demonstrated in a vignette at https://github.com/CshlSiepelLab/DENR/blob/master/vignettes/introduction.Rmd.

## Data availability

All published datasets were downloaded from GEO. GRO-cap data for TSS detection model training was retrieved in preprocessed form using accession number GSE60456 (23). PRO-seq data from K562 (25) and CD4^+^ T cells (33) were retrieved using accession numbers GSE96869 and GSE85337. Data submission of newly generated PRO-seq and RNA-seq to dbGaP is in progress (project #phs002146.v1.p1).

## Acknowledgements

We thank Lingjie Liu for help with data visualization, and other members of the Siepel and Danko laboratories for helpful discussions. This research was supported, in part, by US National Institutes of Health grants R35-GM127070 and R01-HG009309, and by the Simons Center for Quantitative Biology at Cold Spring Harbor Laboratory. The content is solely the responsibility of the authors and does not necessarily represent the official views of the US National Institutes of Health.

