## supplement for "Deconvolution of Expression for Nascent RNA Sequencing Data (DENR) Highlights Pre-RNA Isoform Diversity in Human Cells"

| Transcript ID | Model name | Abundance |
|---|---|---|
| ENST00000393446<br>ENST00000393451<br>ENST00000323984<br>ENST00000393449<br>ENST00000446490 | G14406M1 | 29.6 |
| ENST00000432298<br>ENST00000422922 | G14406M6 | 42.9 |

**Table S1. DENR isoform-abundance estimates for *ST7***

| Gene name | Transcript ID | Model name | Abundance |
|---|---|---|---|
| SEC22C | ENST00000423701<br>ENST00000273156<br>ENST00000449617<br>ENST00000264454 | G10933M2 | 60.5 |
| | ENST00000450981 | G10933M6 | 8.3 |
| SS18L2 | ENST00000447630 | G10431M1 | 30.6 |
| | ENST00000011691<br>ENST00000474941 | G10431M2 | 70.3 |
| NKTR | ENST00000232978<br>ENST00000429888<br>ENST00000617821 | G10432M1 | 132.1 |
| | ENST00000460910 | G10432M3 | 4.6 |
| | ENST00000459950 | G10432M5 | 29.1 |

**Table S2. DENR isoform-abundance estimates for *SEC22C*, *SS18L2* and *NKTR***

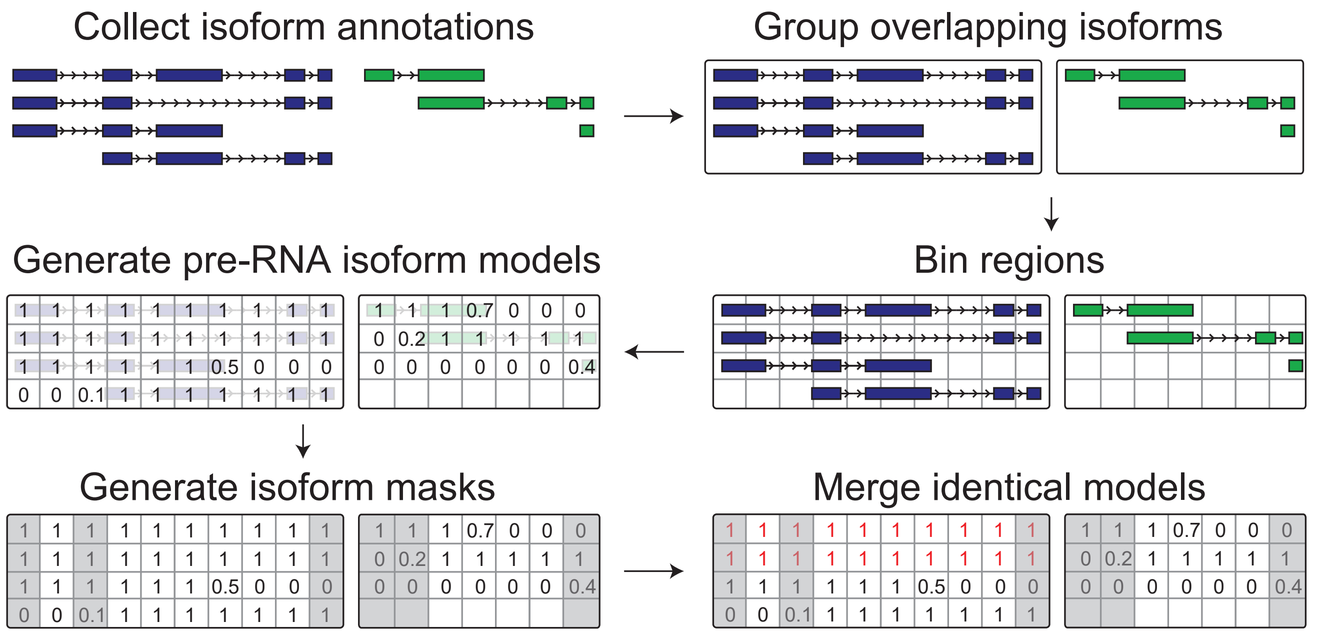


**Fig. S1. Processing isoform annotations for DENR.** This workflow shows the steps involved in generating pre-RNA isoform models from mature RNA isoform annotations. Mature RNA isoform annotations were first downloaded from a database, then grouped into clusters. Bins were overlaid on each cluster (default bin size: 250bp), and presence or absence of annotations in each of these bins was recorded in a design matrix. A user-selected number of bins were optionally masked at the start and end of each isoform. Finally, identical pre-RNA isoforms after masking were merged.

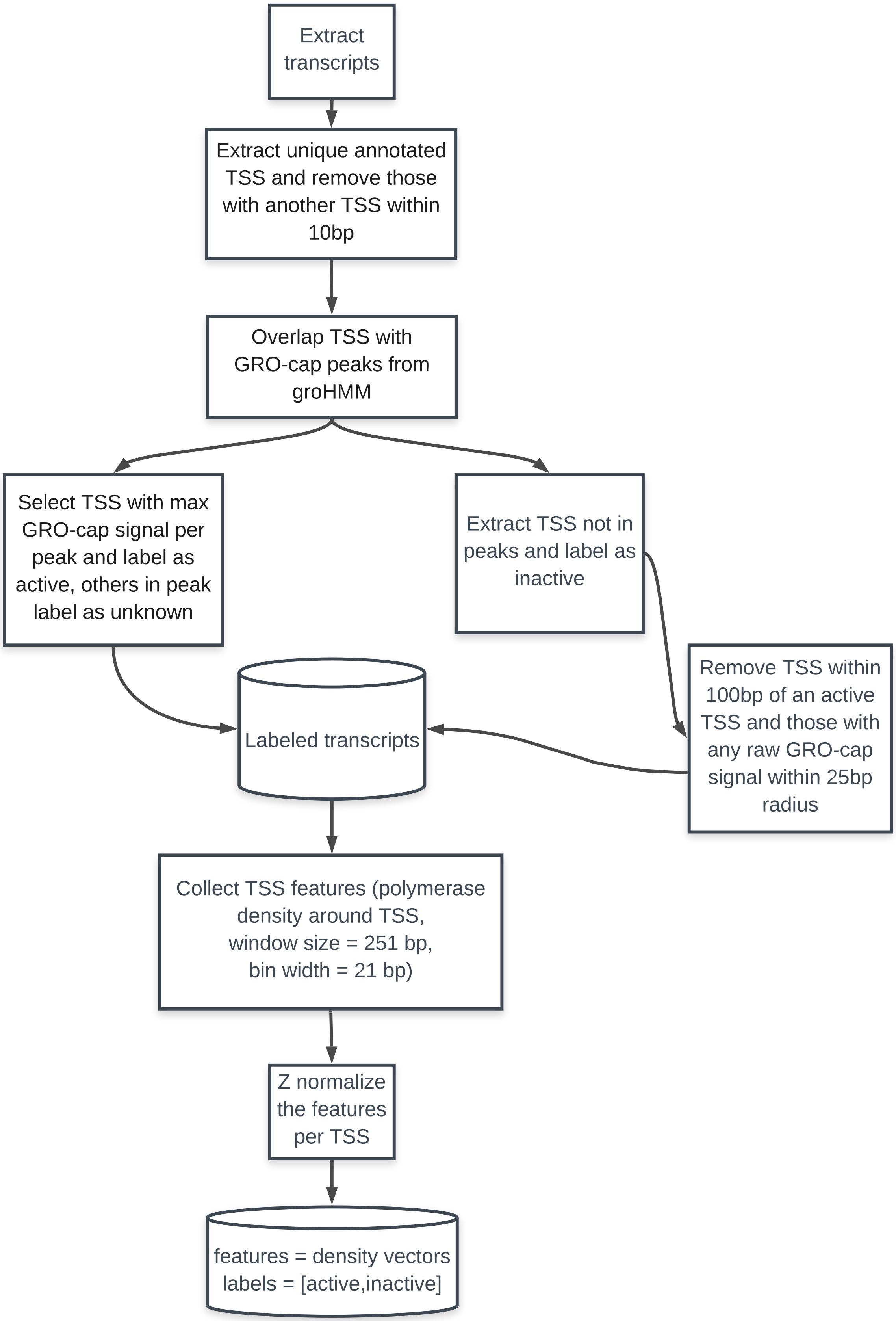


**Fig. S2. Workflow for annotating training data for the TSS classifier**

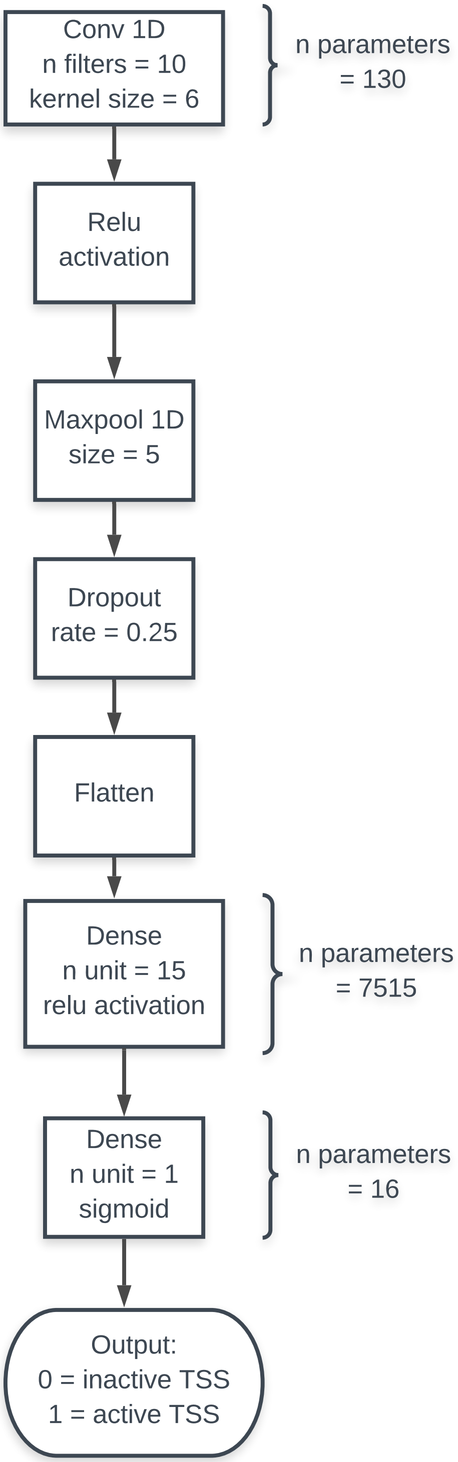


**Fig. S3. Architecture of convolutional neural network for identifying active TSSs**

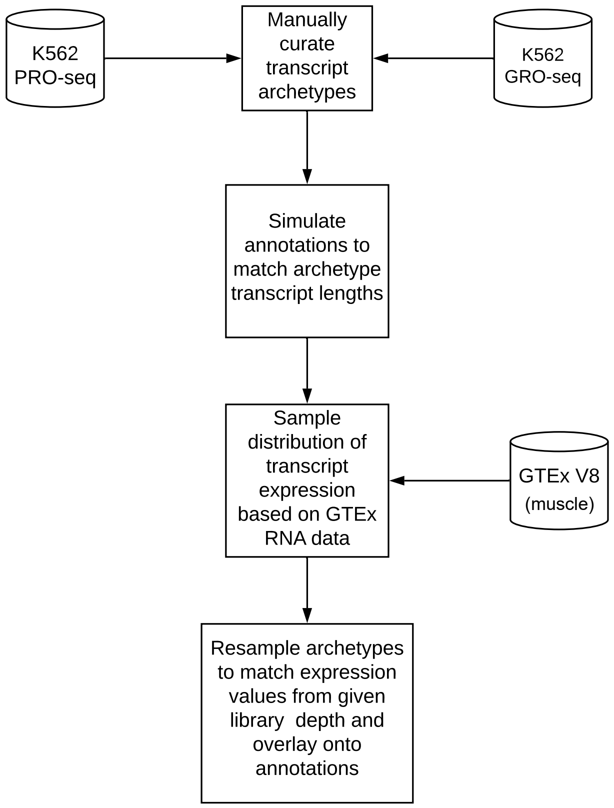


**Fig. S4. Algorithm for empirically simulating nascent RNA sequencing data.**

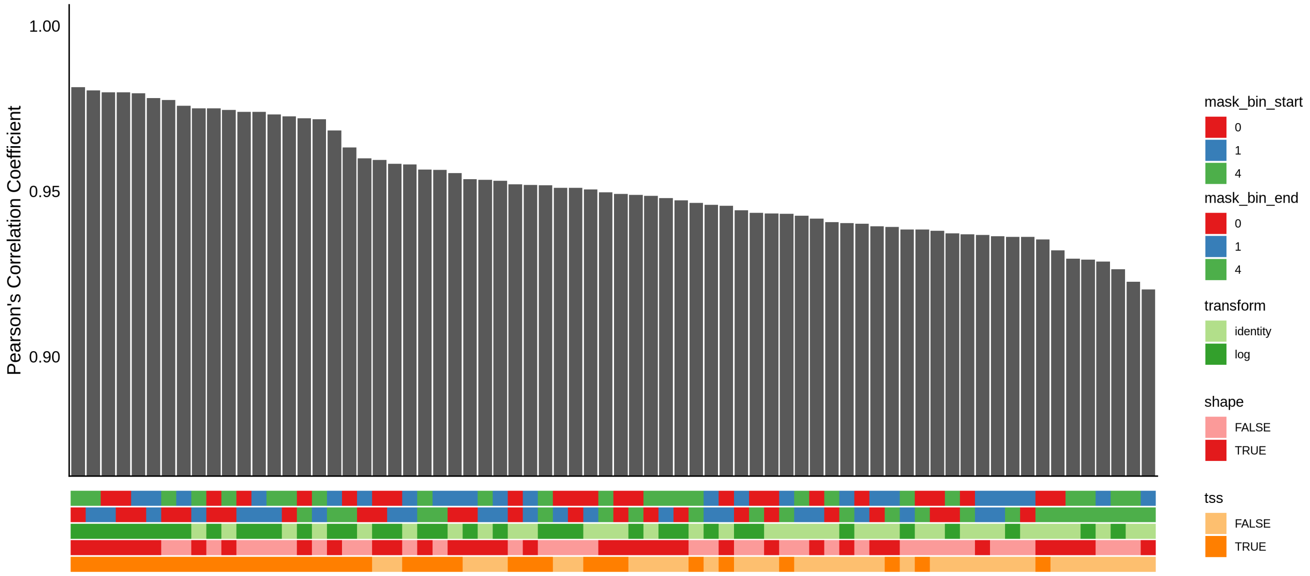


**Fig. S5. Benchmarking of DENR with various combinations of optional features (gene-level).** DENR was run with various numbers (0, 1, or 4) of bins masked at the TSS and TTS of each isoform, and with or without log-transformation of read-counts, shape-profile correction, and TSS prediction ($3 \cdot 3 \cdot 2 \cdot 2 \cdot 2 = 72$ combinations, indicated on the $x$ axis). Bars represent Pearson's correlation coefficient ($r$) between DENR estimates and true values on the gene level.

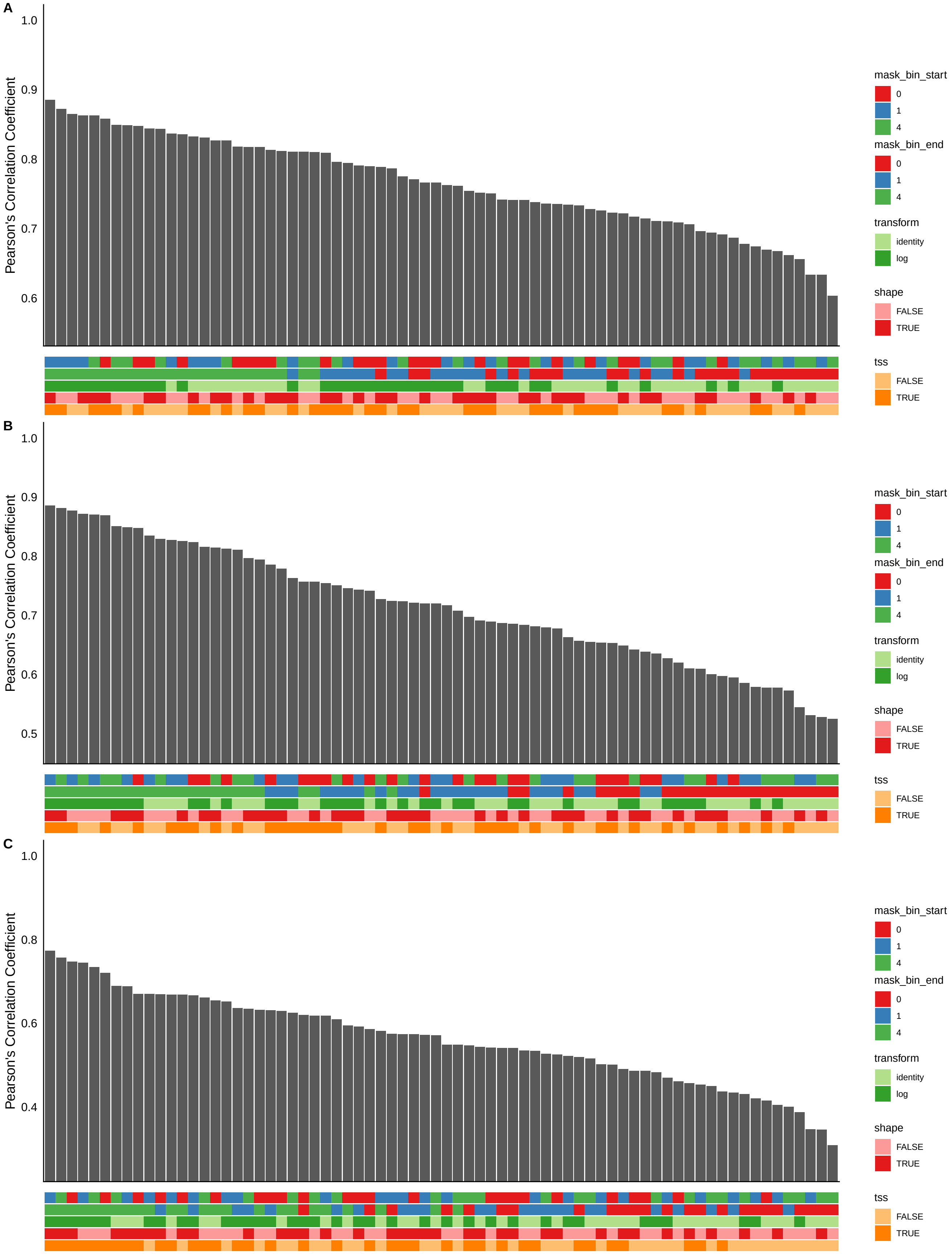


**Fig. S6. Benchmarking of DENR with various combinations of optional features (isoform-level).** DENR was run with the same 72 combinations of optional features as in **Fig. S5**. Bars represent Pearson's correlation coefficient $(r)$ between DENR estimates and true values on the isoform level. **(A)** Dominant isoform. **(B)** Longest isoform. **(C)** All isoforms.

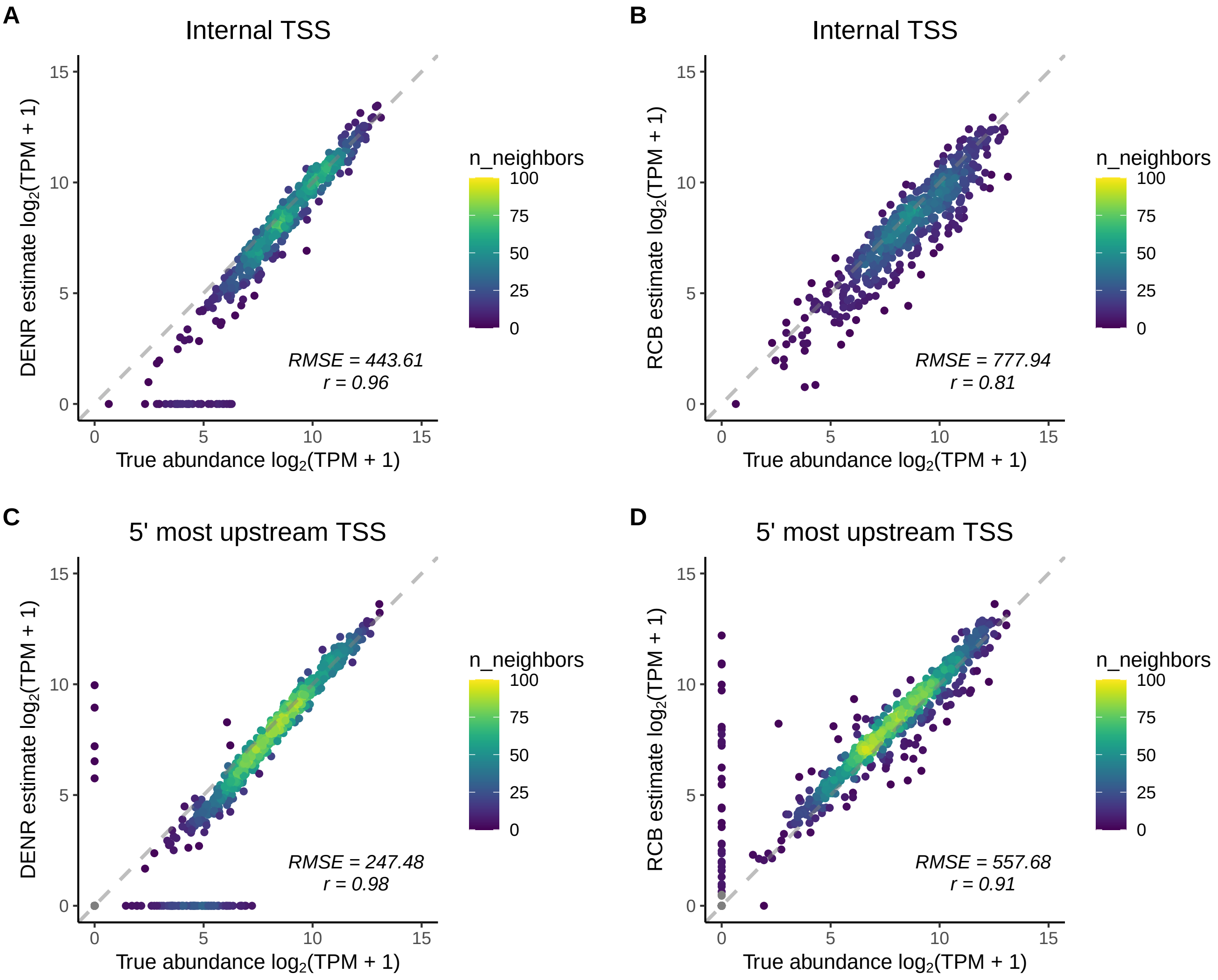


**Fig. S7. Comparison of DENR and the RCB method on subsets of genes of interest. (A)** DENR estimates for genes whose dominant isoform corresponds to an internal TSS. **(B)** RCB estimates for genes whose dominant isoform corresponds to an internal TSS. **(C)** DENR estimates for genes whose dominant isoform corresponds to the 5′-most upstream TSS. **(D)** RCB estimates for genes whose dominant isoform corresponds to the 5′-most upstream TSS. RMSE = root-mean-square error, $r$ = Pearson's correlation coefficient.

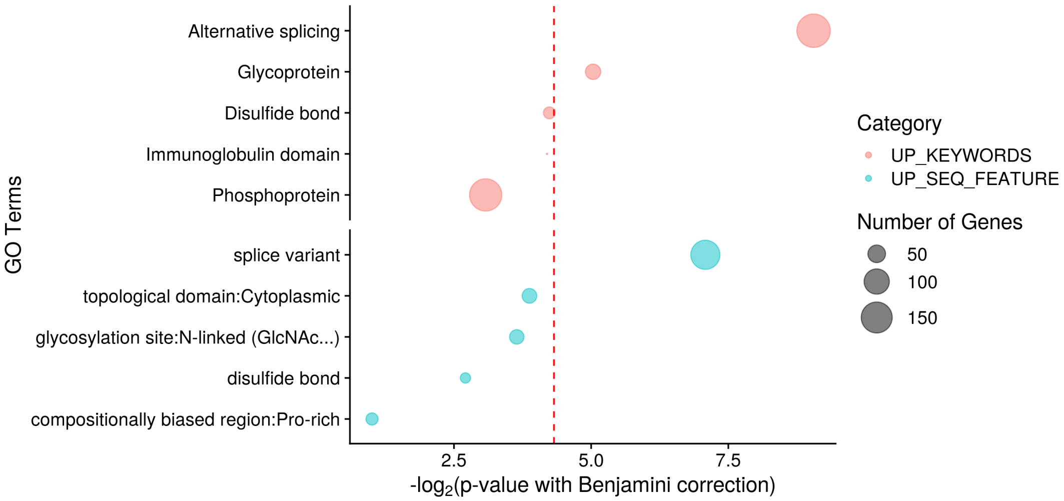


**Fig. S8. Gene Ontology analysis for genes using different TSSs in K562 and CD4+ T cells**. Gene Ontology enrichment for UniProt keywords and sequence annotations with top 5 smallest *p-values* were shown on y axis.

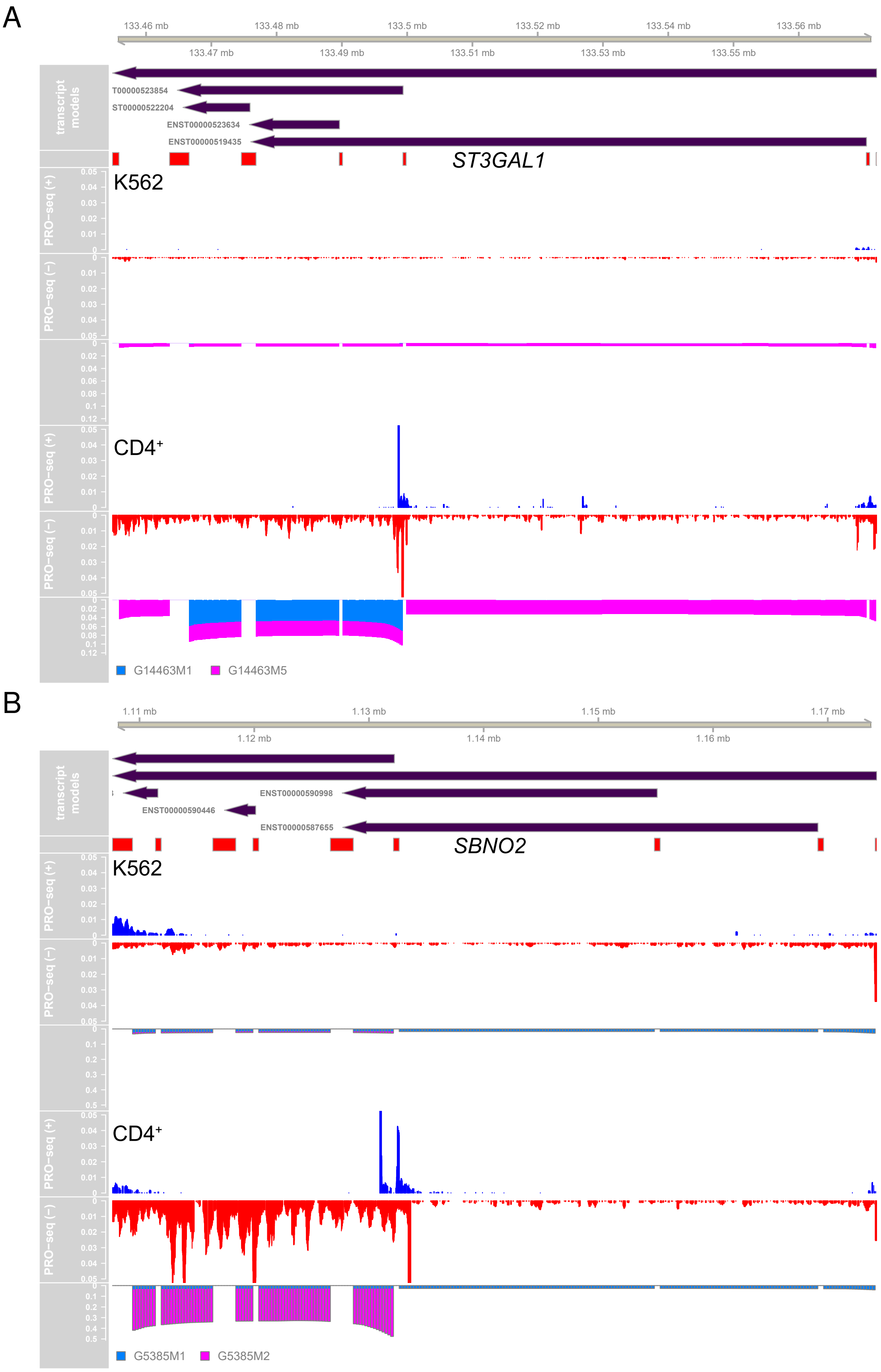


**Fig. S9. Cell type specific TSS usage.** Additional examples for genes using different TSSs in K562 and CD4+ T cells. **(A)** *ST3GAL1* **(B)** *SBNO2*

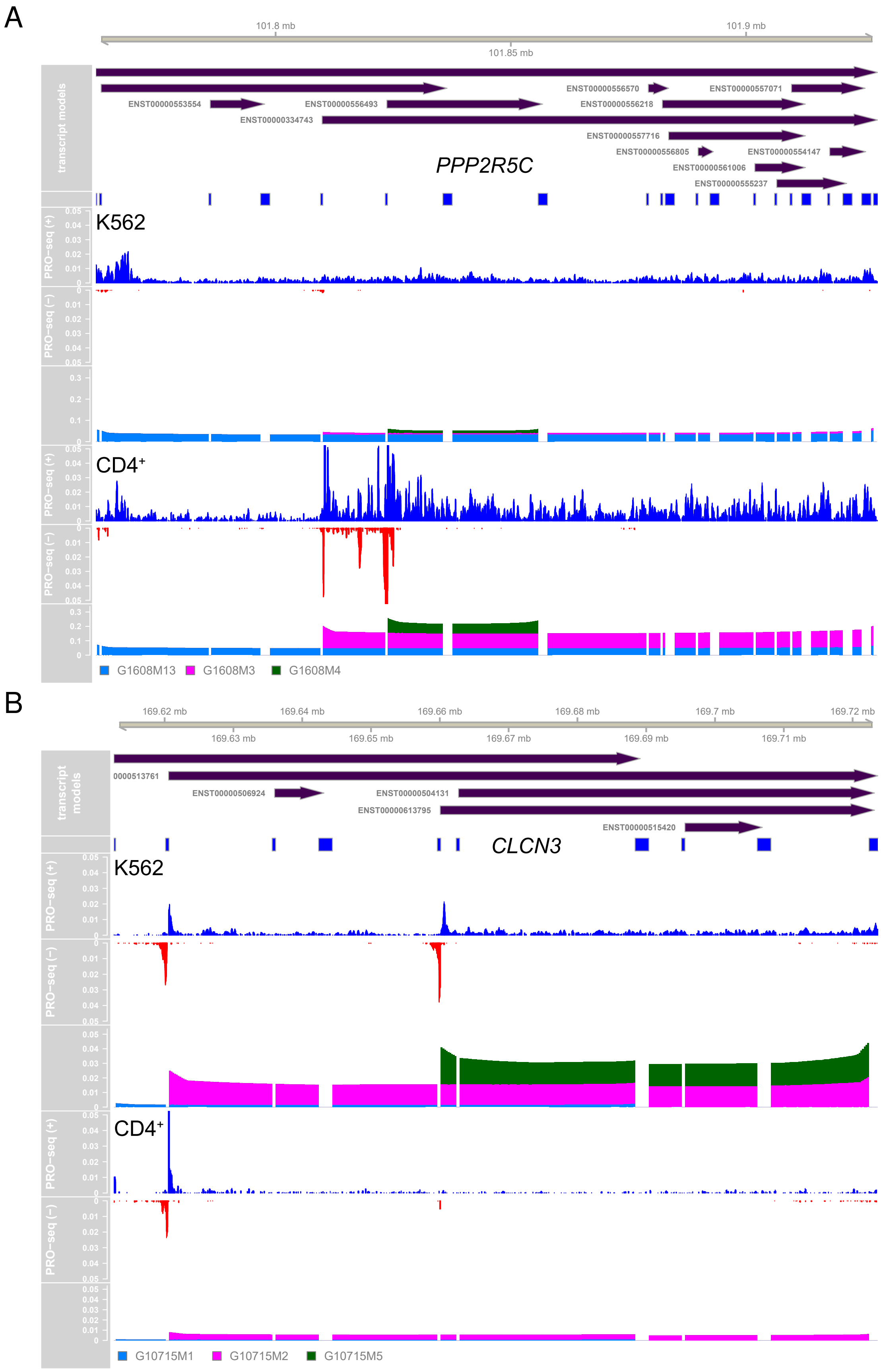


**Fig. S10. Cell type specific TSS usage.** Additional examples for genes using different TSSs in K562 and CD4+ T cells. **(A)** *PPP2R5C* **(B)** *CLCN3*
